# TRPS1 maintains luminal progenitors in the mammary gland by repressing SRF/MRTF activity

**DOI:** 10.1101/2023.11.24.568520

**Authors:** Marie Tollot-Wegner, Marco Jessen, KyungMok Kim, Adrián Sanz-Moreno, Nadine Spielmann, Valerie Gailus-Durner, Helmut Fuchs, Martin Hrabe de Angelis, Björn von Eyss

## Abstract

The transcription factor TRPS1 is a context-dependent oncogene in breast cancer [1] [2] [3] [4] [5]. In the mammary gland, TRPS1 activity is restricted to the luminal population and is critical during puberty and pregnancy [2]. Its function in the resting state remains however unclear. To evaluate whether it could be a target for cancer therapy, we investigated TRPS1 function in the healthy adult mammary gland using a conditional ubiquitous depletion mouse model where long-term depletion does not affect fitness. We show that TRPS1 activity is essential to maintain a functional luminal progenitor compartment. This requires the repression of both YAP/TAZ and SRF/MRTF activities, TRPS1 represses SRF/MRTF activity indirectly by modulating RhoA activity. Our work uncovers a hitherto undisclosed function of TRPS1 in luminal progenitors intrinsically linked to mechanotransduction in the mammary gland. It also provides new insights into the oncogenic functions of TRPS1 as luminal progenitors are likely the cells of origin of many breast cancers.

**Significance statement:** The transcription factor TRPS1 is a context-dependent oncogene in breast cancer. It is unclear how TRPS1 contributes to cancer development and whether it could be a target for therapy. Here we established a mouse model mimicking the systemic effect of TRPS1 drug targeting. With this model, we can show that TRPS1 depletion does not impact the fitness of the animals and that the role of TRPS1 is to maintain a functional luminal progenitor pool in the mammary gland. Mechanistically, TRPS1 represses a mechano-transduction program preventing their commitment to an alveolar fate. Because there is growing evidence that breast cancer originates from the expansion of altered luminal progenitors, our work provides valuable insights into the understanding of breast cancer initiation.

## Introduction

Tricho-Rhino-Phalangeal Syndrome 1 (TRPS1), the only repressor in the GATA family of transcription factors, was originally identified as a regulator of skeletal and cartilage development. Mutations in the *TRPS1* gene cause the autosomal dominant TRP syndrome characterized by skeletal and cranofacial anomalies that are also observed in mice mutants [6]. *Trps1* knockout mice or *Trps1* loss of function mutants lacking the GATA domain die shortly after birth due to respiratory distress caused by malformations of the rib cage [7, 8]. Recently additional functions for TRPS1 have been uncovered in the development of the mammary gland. There, TRPS1 is required for the initial branching of the mammary ducts at puberty but also plays a crucial role in the lactogenic differentiation during pregnancy [9].

It is now well established that TRPS1 can act as a context-dependent oncogene in several breast cancer subtypes [3, 4, 9, 10]. In cancer cells, TRPS1 recruits repressor complexes to enhancer sites which is critical for cancer cell proliferation [3, 11]. In particular, TRPS1 represses the activity of the Hippo pathway transcriptional co-activator Yes-associated protein (YAP) to promote immune evasion [3]. TRPS1 also modulates ER binding at enhancer sites to promote ER-dependent cell proliferation [11]. The *TRPS1* gene is amplified in about 28% of breast tumors which is associated with poor survival prognosis [3]. TRPS1 activity is particularly high in Luminal B and Basal-like breast cancers which are the most aggressive subtypes [3, 5]. In contrast, in Invasive Lobular Carcinoma (10-15% of BC), TRPS1 can also act as a tumor suppressor, in this case it’s the loss of TRPS1 that, associated to loss of E-Cadherin, drives tumor growth [9]. Therefore, understanding the molecular circuits that determine whether TRPS1 has pro- or anti-oncogenic properties is an important issue.

Breast cancer originates in the mammary gland epithelium, which is composed of two cellular lineages, Keratin8-positive (Krt8^pos^) luminal cells and Keratin14-positive (Krt14^pos^) myoepithelial cells forming respectively the inner and outer layer of the mammary ducts. During lactation, the luminal lineage gives rise to milk secretory cells while the contractile basal cells enable milk expulsion. An expansion of aberrant luminal progenitors expressing basal markers is induced by aging or at a young age in BRCA1^mut^ and BRCA2^mut^ carriers [12]. These aberrant progenitors are believed to be the cells of origin of most breast cancer types, and their expansion dramatically increases the risk to develop breast cancer [12–16].

Various regulators have been shown to control lineage commitment and differentiation in the mammary epithelium. Among them, the hippo transducer TAZ promotes the commitment to the basal lineage. Forced expression of TAZ in luminal cells induces them to adopt basal characteristics and depletion of TAZ in basal cells leads to luminal differentiation [17]. In the luminal lineage Elf5 is exclusively expressed in the luminal progenitors of the resting mammary gland and induces the differentiation into milk-secretory cells during pregnancy. Stat5 is specifically involved in the maintenance and regeneration of the luminal progenitors [18] and Gata3 promotes their differentiation into luminal alveolar cells [19, 20]. Similarly, Trps1 activity is restricted to the luminal population [9] besides its critical role in lactation, the function it fulfils in the healthy resting mammary gland has not yet been studied. Yet this knowledge would be essential to evaluate whether it could be a future target for cancer therapy.

With this in mind, we set out to investigate the function of TRPS1 in the homeostasis of the adult mammary epithelium. Using a ubiquitous depletion mouse model where long-term depletion does not affect fitness, we show that TRPS1 activity is essential to maintain a functional luminal progenitor population. Besides the direct repression of YAP/TAZ targets, TRPS1 is required to repress an SRF/MRTF transcriptional program since elevated activity of the mechanosensitive SRF/MRTF circuitry is sufficient to inhibit progenitor function. This repression is indirect and involves a set of TRPS1-regulated genes encoding RhoA modulators. Our work uncovers a hitherto function of TRPS1 in luminal progenitors which is intrinsically linked to mechanotransduction in the mammary gland. It also provides new insights into the oncogenic functions of TRPS1.

### TRPS1 long-term depletion has no deleterious effect in adult female mice

To systematically analyze whether TRPS1 could potentially be used as a target for cancer therapy, it is vital to understand its role for organismal fitness in the whole body. To this end, we generated a mouse line where we could study the effects of TRPS1 depletion *in vivo* by depleting it in the whole mouse. Since *Trps1* KO mice are not viable we used a model where TRPS1 could be depleted in the adult animal in order to ensure proper embryonic development. In addition, we wanted the depletion to be effective in the whole body to mimic the systemic effect of TRPS1 drug targeting. To fulfill these different criteria, we chose to deplete TRPS1 using ubiquitous doxycycline-inducible shRNA expression and generated a mouse line harboring a TRE-turboGFP-shRNA cassette in the *Col1a1* locus and a CAG-rtTA3 transgene (Fig 1a, S1a). With this model, TRPS1 can be depleted at any time by giving the animals doxycycline via food. The cells expressing the shRNAs can then be identified based on turboGFP (tGFP) expression (Fig 1a). We generated two distinct lines expressing different Trps1 shRNAs, as well as a control line targeting Renilla and verified the efficiency of each Trps1 shRNA in MEFs isolated from transgenic animals (Fig 1b).

**Figure 1:**
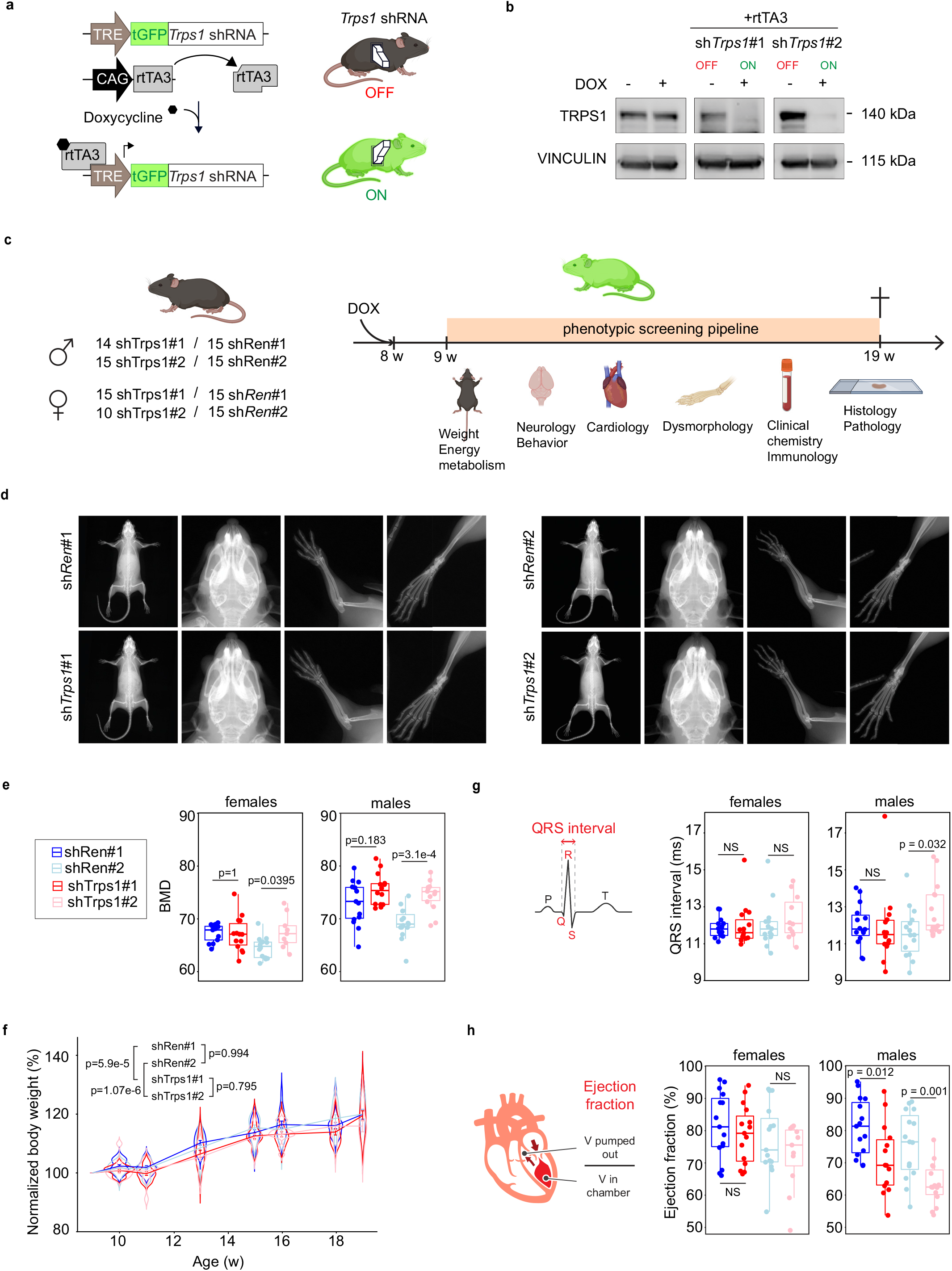
Ubiquitous TRPS1 depletion does not affect fitness. (a) Overview of the genetic structure of the ubiquitous TRPS1 depletion model mouse line. The line harbors a CAG-rtTA3 transgene composed of CAG promoter followed the reverse tetracycline-controlled transactivator 3 (rtTA3) sequence. Sequences allowing the expression of a tGFP marker and Trps1 shRNA under the control of Tetracyclin Response Element (TRE) are inserted in the *Col1a1* locus. Their expression is enabled by rtTA3 in the presence of doxycycline. (b) Immunoblot of mouse embryonic fibroblasts isolated from TRPS1-depletion lines showing the depletion of TRPS1 upon Dox-induced shRNA expression. Vinculin serves as a loading control. (c) Schematic of the phenotypic screening performed at the german mouse clinic. Animals used for the screening (left); males and females from shTrps1 #1 and #2 depletion lines and shRen #1 and #2 corresponding littermate controls. Right panel: short description of the screening pipeline. (d) X-rays pictures of the whole body, skull, arm, and leg from TRPS1 depletion mice and shRenilla littermate control mice. (e) Box plot of the bone marrow density of male and female animals of the indicated mouse lines. Females: shRen#1 n=15, shTrps1#1 n=14, shRen#2 n=14, shTrps1#2 n=9. Males: shRen#1 n=15, shTrps1#1 n=14, shRen#2 n=15, shTrps1#2 n=14. One-way ANOVA with Tukey HSD, post-hoc test. (f) Body weight of female animals from the indicated mouse lines during the 10 week-phenotypic screen. shRen#1 n=15, shTrps1#1 n=15, shRen#2 n=15, shTrps1#2 n=11. Two-way ANOVA, with Tukey HSD post-hoc test. (g) QRS interval (ms) of male and female animals from each mouse line. Females: shRen#1 n=15, shTrps1#1 n=15, shRen#2 n=15, shTrps1#2 n=11. Males: shRen#1 n=15, shTrps1#1 n=14, shRen#2 n=15, shTrps1#2 n=15. Wilcoxon rank sum test. (h) Ejection fraction (%) of male and female animals from each mouse line. Females: shRen#1 n=15, shTrps1#1 n=15, shRen#2 n=15, shTrps1#2 n=11. Males: shRen#1 n=15, shTrps1#1 n=14, shRen#2 n=15, shTrps1#2 n=15. Wilcoxon rank sum test.

To evaluate the long-term effects of *Trps1* depletion, we subjected our mice to a detailed phenotypic screening over a period of ten weeks (from nine weeks to nineteen weeks of age). About 15 mice for each sh*Trps1* line were screened together with their respective shRen control litter mates (Fig 1c, S1a). The phenotyping pipeline included various behavioral and neurological tests, a dysmorphology analysis, energy metabolism and cardiac function evaluations, as well as clinical chemistry and immunology screenings (the full screening pipeline is described in Figure S2). During the whole screening time, the mice were kept on doxycycline, and their body weight was measured weekly. Because TRPS1 is required for embryonic skeletal and cartilage development, we paid particular attention to the dysmorphology analysis. Full body X-Rays imaging did not reveal genotype-specific anatomical alterations (Fig 1d). By DXA, bone mineral density (BMD) was slightly increased in the shTrps1#2 line when compared to the respective shRen#2 controls, especially males (Fig 1e, S2). These differences were not as clear with the shTrps1#1 construct (Fig 1e). Only minor differences in body weight were observed between shTrps1 and control genotypes (Fig. 1f). At the end of the phenotyping pipeline, the animals were sacrificed, and histological analysis carried out on 33 organs/tissues (n=5 per genotype/sex). No obvious genotype-specific histopathological alterations were found between animals depleted for TRPS1 and controls, including in the mammary gland tissue. Males of both sh*Trps1* lines also showed altered cardiac function with a lower ejection fraction for both lines and a longer QRS interval for sh*Trps1*#2 animals (Fig. 1g and 1h). Apart from the minor alterations observed in males, TRPS1 depletion did not cause any significant deleterious effect in female mice. This suggests that systematic TRPS1 inhibition could be a valid therapeutic strategy.

### TRPS1 regulates luminal progenitor gene expression in the homeostatic mammary gland

In the mammary gland epithelium, luminal cells depend on TRPS1 activity for survival and differentiation [9]. We examined the morphology of the mammary ducts after TRPS1 depletion. Here, we did not observe any difference between sh*Trps1* and sh*Ren* animals treated with doxycycline (Fig 2a), even though TRPS1 was efficiently depleted in vivo (Fig 2b). However, the mammary gland is a highly plastic tissue and TRPS1-negative cells are rapidly counter-selected during mammary gland development [9], suggesting that TRPS1-dependent phenotypes may be masked by compensatory mechanisms.

**Figure 2:**
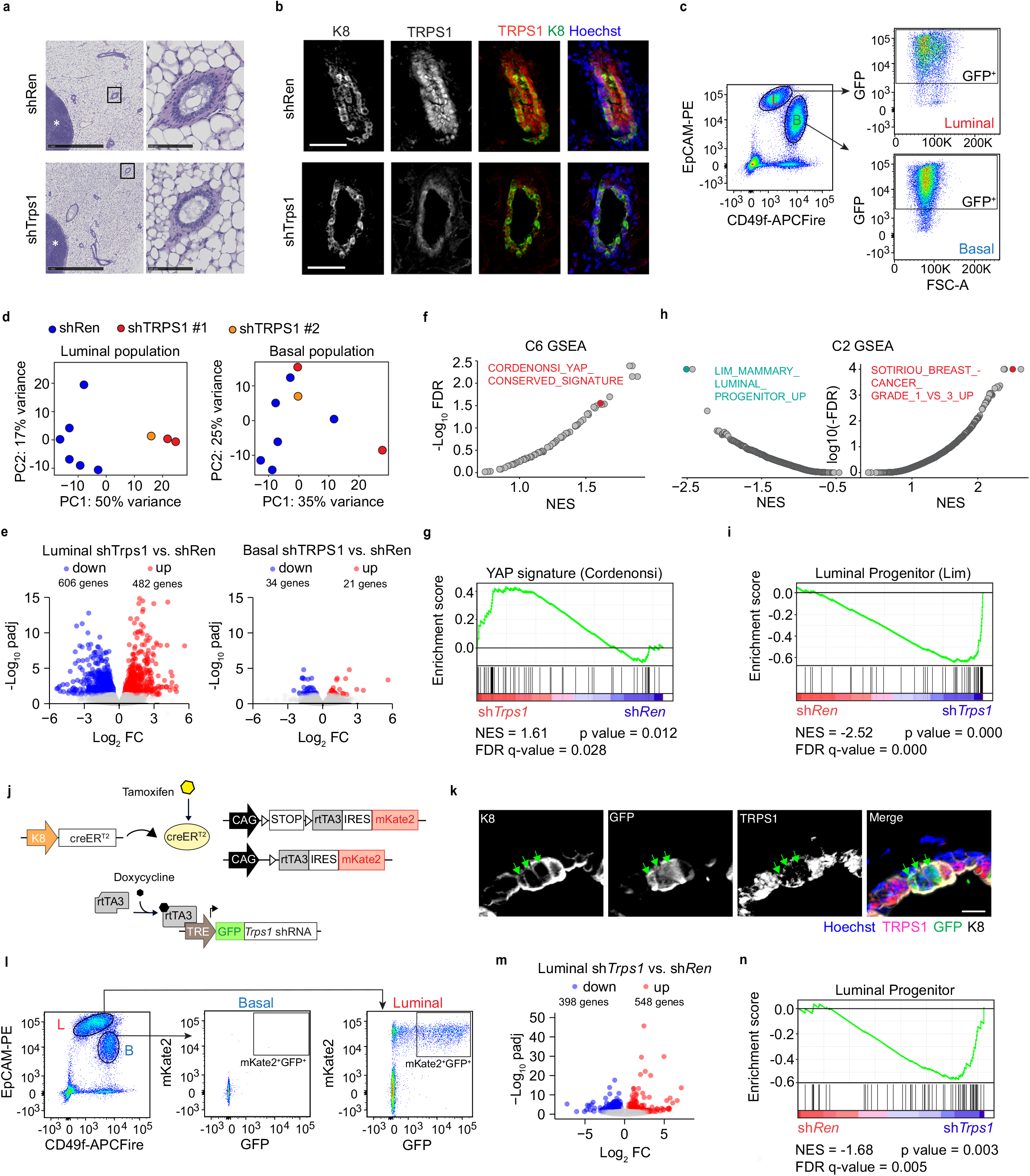
TRPS1 controls a luminal progenitor transcriptional program. (a) H & E stainings of mammary gland cross-sections from shRen and shTrps1 mice treated with Doxycycline for 10 weeks. Left: overview. Right: close up of a mammary duct. scale bars = 50 um. (b) Immunofluorescent stainings of the indicated proteins in mammary ducts from shRen and shTrps1 female mice treated for 10 days with doxycycline. Scale bar = 50 um. (c) Representative flow cytometry plots of Luminal and basal cells expressing shRNAs (GFP^+^) (d) Principle Component Analysis (PCA) plots RNA-Seq data from Luminal and basal cells expressing shRen or shTrps1 (e) Volcano plots showing differential gene expression in luminal (left) and basal (right) cells expressing shTrps1 vs shRen. (f) Gene set enrichment analysis (GSEA) of shTrps1 vs shRen luminal cells using C6 oncogenic gene sets. (g) GSEA Enrichment plot for YAP signature (Cordenonsi et al., 2011) in shTrps1 vs shRen luminal cells. (h) Gene set enrichment analysis of shTrps1 vs shRen luminal cells using C2 curated gene sets. (i) GSEA Enrichment plot for Luminal progenitor signature (Lim et al., 2009) signature in shTrps1 vs shRen luminal cells. (j) Overview of the genetic structure of the luminal-specific TRPS1 depletion mouse line. The line harbors two transgenes. One is composed of the Krt8 promoter and the CreERT2 recombinase coding sequence. The other contains rtTA3 and mKate2 coding sequences under the control of a CAG promoter. A Lox-STOP-Lox prevents their expression. In addition a TRE-tGFP-shRNA element is inserted in the *Col1a1* locus. Tamoxifen allows excision of STOP by recombinase and expression of rtTA3 and mKate2. Expression of tGFP and shRNA is enabled by rtTA3 in the presence of doxycycline. (k) Immunofluorescent stainings of the indicated proteins in mammary ducts from luminal-specific TRPS1-depletion mouse. Scale bar = um. (l)) Representative flow cytometry plots of recombined (mKate2^+^) Luminal and basal cells expressing shRNAs (GFP^+^) (m) Volcano plots showing differential gene expression in luminal cells expressing shTrps1 vs shRen (n) GSEA Enrichment plot for Luminal progenitor signature (Lim et al., 2009) in shTrps1 vs shRen luminal cells

For this reason, we investigated how TRPS1 influences the mammary gland transcriptome. We isolated tGFP-positive (shRNA expressing) luminal and basal cells (Fig. 2c and S4) from mice kept for 28 days on doxycycline and performed bulk RNA-Sequencing. The luminal samples showed clear separation in the Principal Component Analysis (PCA) using the 1000 most variable genes (Fig. 2c,d). In contrast only a handful of genes showed differential expression between basal sh*Ren* and sh*Trps1*, consistent with the fact that TRPS1 expression is restricted to luminal cells (Fig. 2b). As in breast cancer cells [3], the expression of YAP target genes was strongly induced by depletion of TRPS1 (Fig. 2f,g). In addition, gene sets associated with low grade breast cancer were upregulated in shTRPS1 luminal cells (LCs) (Fig. 2h) which points to the fact that TRPS1 can be an oncogene in breast cancer.

Interestingly, genes associated with luminal progenitor function [12] were most significantly downregulated in shTrps1 luminal cells (Fig. 2h,i). We next validated the cell-intrinsic nature of luminal TRPS1 depletion on the luminal progenitor program, by using a K8-CreER mouse line to restrict shRNA expression to luminal cells (Fig. 2j-l, and S5). TRPS1 depletion in this model also led to profound changes in the luminal cells (Fig. 2m), and the same luminal progenitor gene set was also downregulated upon TRPS1 depletion (Fig. 2n). Collectively, these results demonstrate that luminal expression of TRPS1 is required for maintenance of a luminal progenitor expression program in the adult mammary gland.

### Trps1 regulates luminal progenitor function

To clearly define the function of TRPS1 in luminal progenitors (LPs), we isolated LPs and mature cells from WT mice using the c-Kit surface marker (Kendrick et al., 2008, Regan et al., 2012) (Fig. 3a and S6). As expected, the mature marker Amphiregulin (*Areg*) showed higher expression in mature cells compared to progenitors while the progenitor marker *Elf5* was expressed at a higher level in the LP fraction (Fig. 3b). *Trps1* expression was approximately five times higher in the LP population compared to mature cells (Fig. 3b). To identify the LP-specific gene expression signature, we performed RNA-Seq on c-Kit^high^ (LP) and c-Kit^low^ (mature) luminal fractions. Differential gene expression comparing the two sorted populations revealed ∼4000 differentially expressed genes (Log2FC < 1, padj < 1e-4), we defined a gene signature for each cell type (Fig. 3c, Table S1 and S2). Both signatures were downregulated in the bulk RNA-Seq experiment where we compared shRen vs. shTrps1 luminal cells (Fig. 3d, see also Fig. 2e,).

**Figure 3:**
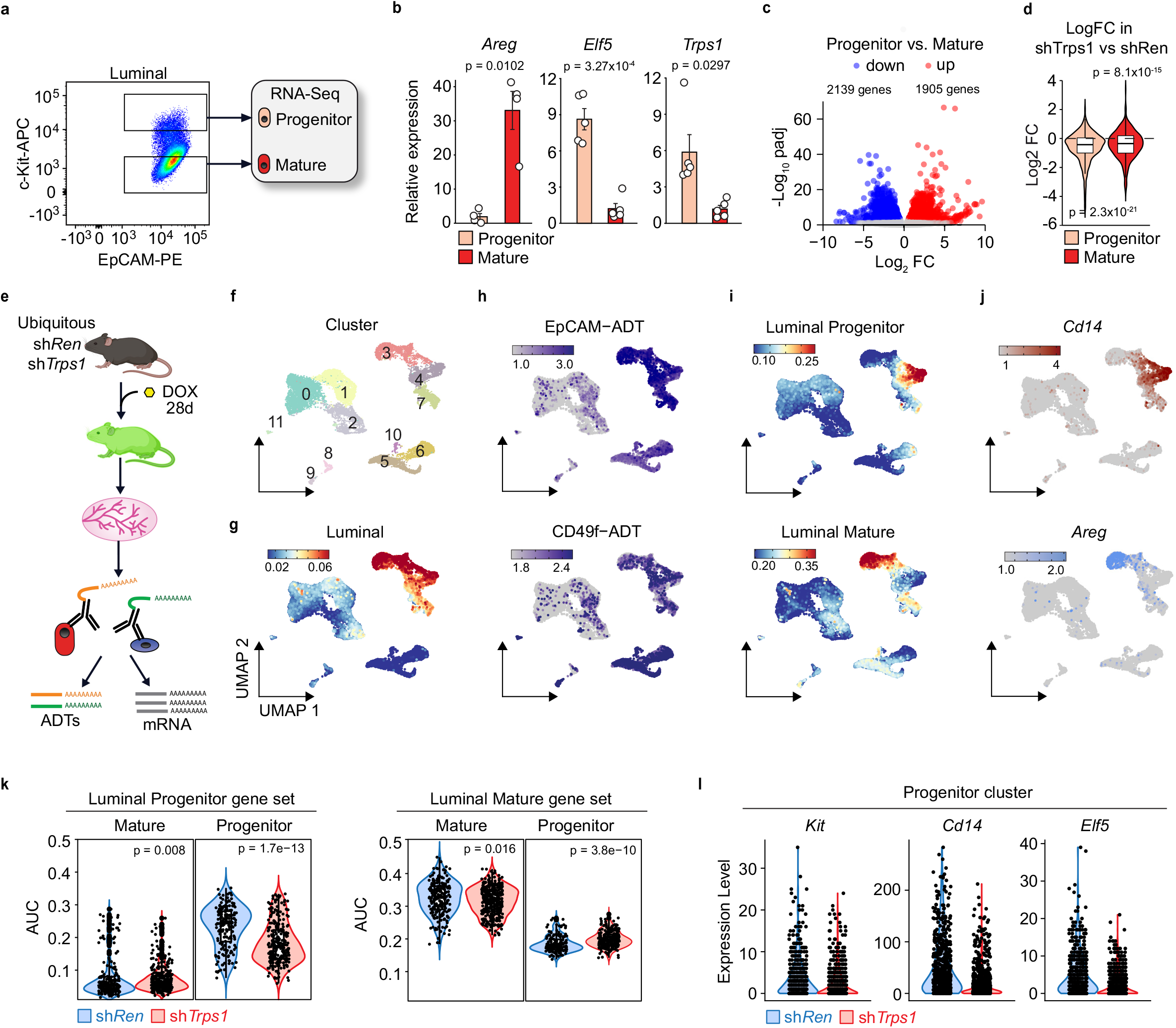
TRPS1 specifically maintains the luminal progenitor population. (a) representative flow cytometry plot of mammary luminal progenitor and mature cells sorted for RNA-Seq based on c-Kit marker. (b) qRT-PCRs of the mature marker gene *Areg*, the progenitor marker gene *Elf5* and *Trps1* in sorted luminal mature and luminal progenitor cells. Summary of 5 biological samples, t-test with Benjamini Hochberg correction, values shown are adjusted p values (p_adj_). (c) Volcano plot of differential gene expression between mammary luminal progenitor and luminal mature cells. padj = adjusted *p*-value, FC = Fold Change. (d) Violin plots for the progenitor and mature gene sets determined in (c) comparing the expression in shTrps1 vs shRen cells (RNA-Seq data Figure 2). (e) Schematic of the CITE-Seq experiment comparing 28 day-dox treated shTrps1 and shRen animals. (f)-(j) UMAP dimensionality reduction plots of the CITE-Seq data from shTrps1 and shRen mammary gland cells combined. Graph-based clusters (f) luminal gene set activity based on AUC (g), EpCAM and CD49f proteins based on ADTs (h), luminal progenitor and luminal mature (determined in c) gene set activity based on AUCs (i), *Cd14* and *Areg* mRNA expression (j) (k) Violin plots of the luminal progenitor (left) and mature (right) gene set activity (c) in mature cluster 3 and progenitor cluster 4 from shTrps1 (red) and shRen (blue) CITE-Seq samples. (l) Expression of *Kit*, *Cd14* and *Elf5* in luminal progenitor cluster 4 in shTrps1 (red) compared to shRen (blue) cells. Boxplots in (d): bottom/top of box: 25^th^/75^th^ percentile, upper whisker: min(max(x), Q3+1.5*IQR), lower whisker: max(min(x), Q1-1.5*IQR), center:median.

To test whether *Trps1* depletion indeed specifically affected the luminal progenitor population, and to be agnostic to surface marker expression, we isolated total mammary gland cells from shTrps1 and shRen transgenic mice and carried out Cellular Indexing of Transcriptomes and Epitopes by Sequencing (CITE-Seq) experiments to analyze the transcriptome and the abundance of chosen surface markers at the single cell level (Fig. 3e). Cell clustering based on the gene expression data defined 12 different clusters (Fig. 3f). Cluster 3, 4 and 7 corresponded to the luminal population based on the activity of a luminal gene set and the abundance of EpCAM and CD49f surface markers (Fig. 3g and 3h). Within the luminal population, Cluster 4 was mainly composed of progenitors and cluster 3 of mature cells based on the activity of the luminal progenitor and mature gene sets that we defined in this study (Fig. 3c) and others that were published previously [21] (Fig. 3i and S3). This correlated with the mRNA level of selected marker genes (Fig. 3j). We then compared the activity of the progenitor gene set in sh*Trps1* and control cells in clusters 3 and 4. *Trps1* depletion led to a clear downregulation of the luminal progenitor signature in the progenitor cluster 4 but not in the mature cluster 3 (Fig. 3k). In comparison, the mature gene set activity was only slightly affected by *Trps1* depletion (Fig. 3k). Selected marker genes followed the same pattern (Fig 3l). We thus concluded that Trps1 is mainly expressed in the LP compartment and depletion therefore specifically affects the LP population, whereas changes in the mature population may be largely secondary in nature.

### Trps1 maintains the functionality of the LPs

Next, we examined whether TRPS1 depletion compromised the functionality of luminal progenitors. In addition to c-Kit, we used here CD14 as another luminal progenitor surface marker in flow cytometry since our scRNA-Seq analysis identified CD14 as highly specific LP marker (Fig. 4a, see also Fig. 3j). We analyzed by flow cytometry the proportion of GFP^hi^ cells that remained in the luminal progenitor population after depletion of Trps1 (Fig. 4a, 4b and S7). Because the CAG-rtA3 driver was unable to induce shRNA expression in all cells with always a small but significant proportion of cells remaining GFP^neg^/GFP^lo^ - these cells could then potentially function as competitor cells for GFP^hi^ cells (Fig. 4b). While there was no significant change in the basal population, the luminal population demonstrated drastic differences upon TRPS1 depletion (Fig. 4c). The fraction of GFP^hi^ cells in the LP compartment was reduced almost by 50% in shTrps1-expressing animals compared to shRen controls. We observed the same trend in the mature subpopulation although to a lesser extent and this was reflected in the total luminal population. The reduced contribution of GFP^hi^ LPs to the overall luminal population suggested that the function of progenitors was perturbed. The gold standard for testing the functionality of progenitor cells is the colony formation assay (CFC). Therefore, we sorted GFP^hi^ and GFP^lo^ luminal cells from doxycycline-treated mice and tested their performance in the CFC assay (Fig. 4d, and S8). No significant difference could be observed in the number of colonies formed by GFP^hi^ or GFP^lo^ cells from shRen control animals. However, GFP^lo^ luminal cells from shTrps1 animals formed many more colonies than their GFP^hi^ counterparts indicating a higher proportion of functional progenitors among the GFP^lo^/GFP^neg^ luminal population (Fig. 4e). Surprisingly, the absolute number of colonies formed by GFP^lo^ cells was higher in shTrps1 animals than in shRen. This difference could be a compensatory mechanism: The GFP^lo^ LPs might expand to fulfill the functions of the functionally impaired GFP^hi^ LPs (Fig. 4e). Taken together these results demonstrate that TRPS1 plays an active role in maintaining LP function.

**Figure 4:**
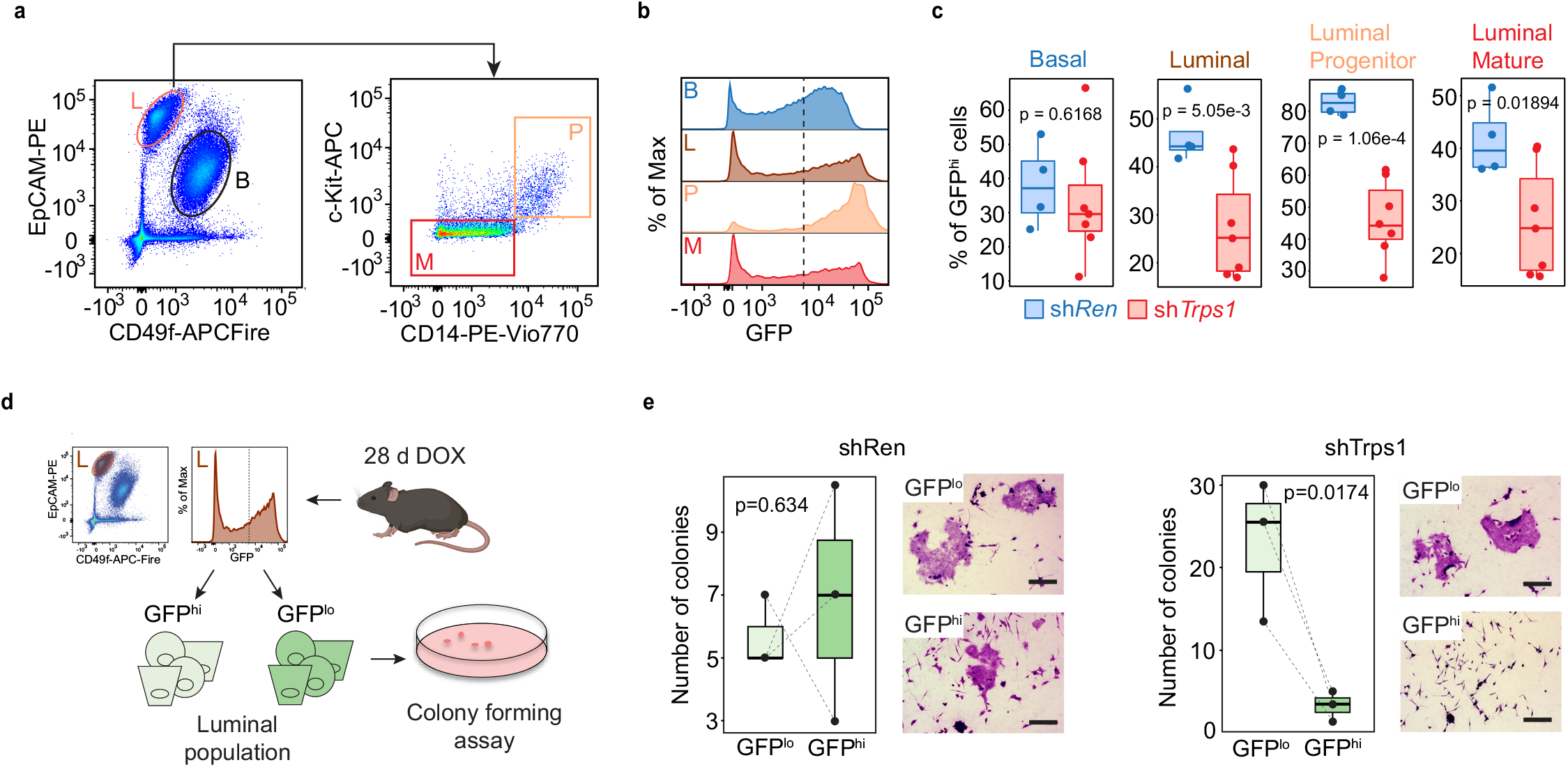
TRPS1 maintains LP function. (a) Representative flow cytometry plots of the different mammary epithelial populations. (b) Representative GFP flow cytometry plot of the different mammary epithelial populations. (c) Proportion of GFP^hi^ cells in the different mammary epithelial populations. shRen, n = 4 animals; shTrps1, n = 7 animals. Welch t-test. (d) Schematic of the colony forming assay using sorted GFP^hi^ and GFP^lo^ luminal cells (e) Number of colonies formed by sorted GFP^hi^ and GFP^lo^ shRen (left panel) and shTrps1 (right panel) luminal cells. Right side: Representative pictures (bar = 300 μm). n = 3 biological samples. One-way ANOVA with Tukey HSD post-hoc test.

### Trps1 represses SRF activity to maintain a functional LP population

To identify TRPS1-regulated genes or pathways that are required to maintain the luminal progenitor function, we went back to our scRNA-Seq data and performed re-clustering of the luminal population. Among the seven newly identified luminal clusters, cluster 2 contained the LP population and showed prominent *Trps1* expression (Fig. 5a). In contrast, mature luminal cells were distributed in clusters 1 and 3 and showed low *Trps1* expression (Fig 5a).

**Figure 5:**
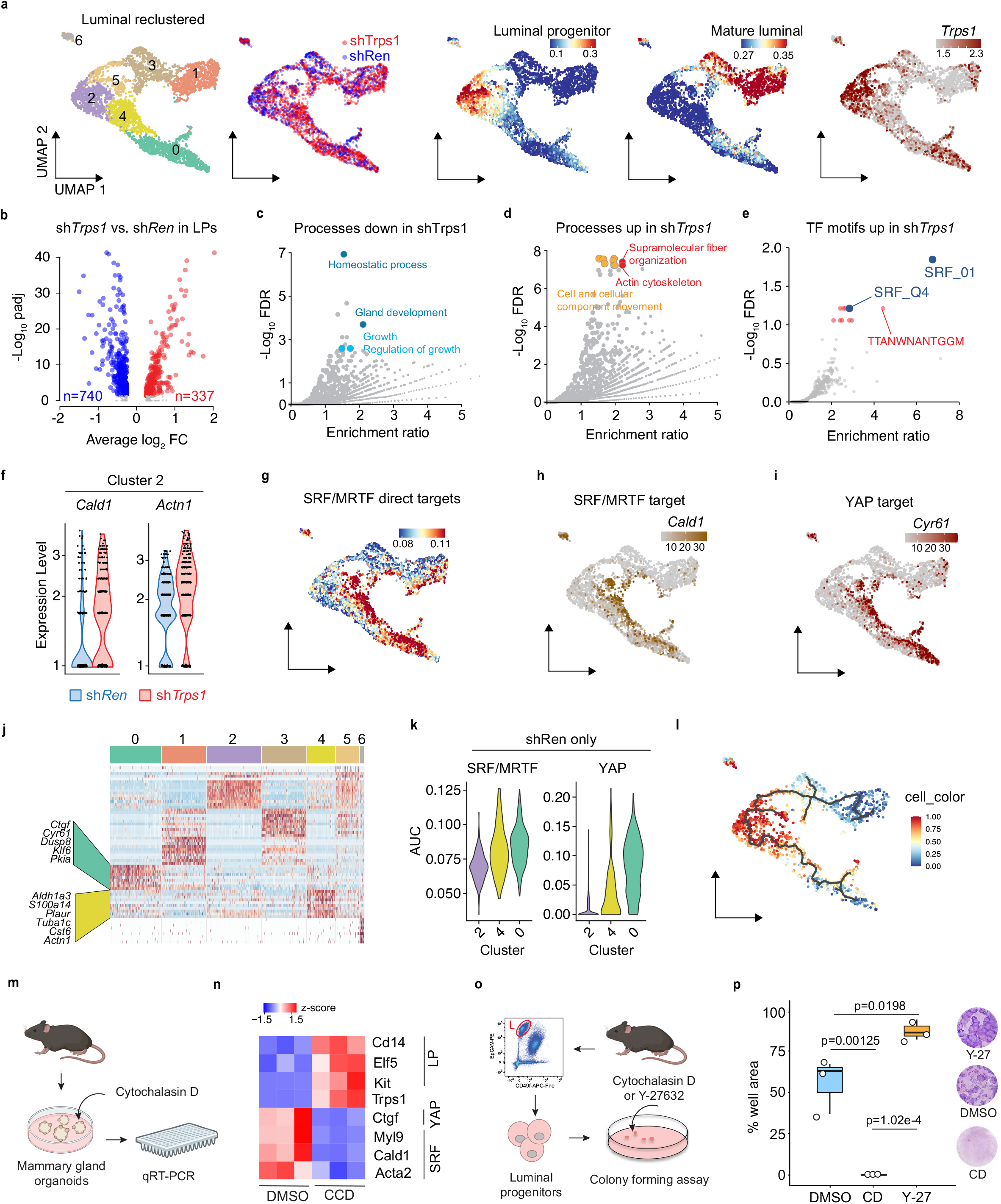
TRPS1 maintains LP function by repressing SRF/MRTF activity. (a) UMAP dimensionality reduction plots of the re-clustered luminal population from CITE-Seq (Figure 3). From left to right: Graph-based clusters, Highlight of shTrps1 (red) and shRen (blue) cells, luminal progenitor and luminal mature gene set (Figure 3c) activity based on AUCs and *Trps1* mRNA expression. (b) Volcano plot of the differential gene expression in shTrps1 vs shRen luminal progenitors (Cluster 2). padj = adjusted *p*-value, FC = Fold Change. (c)-(e) Volcano plots of processes enriched in genes down regulated (c), processes enriched in genes up regulated (d) and TF motifs enriched in genes up regulated in shTrps1 vs shRen luminal progenitor cells (e). FDR = False Discovery Rate. (f) Expression of SRF/MRTF target genes *Cald1* and *Actn1* in luminal progenitor cluster 2 in shTrps1 (red) and shRen (blue). (g) UMAP dimensionality reduction plot of SRF/MRTF gene set activity based on AUCs. (h)(i) UMAP dimensionality reduction plot of *Cald1* (h) and *Cyr61* (i) mRNA expression. (j) Heatmap of the differentially expressed genes in each cluster of shRen control cells. (k) Violin plots of SRF/MRTF (left) and YAP (right)-targets gene set activities in the indicated clusters. (l) CytoTRACE pseudotime trajectory of luminal cells in shRen control animals (m) Experimental design: SRF was activated in mammary organoids by Cytochalasin D treatment and gene expression changes evaluated by qRT-PCR. (n) Heatmap based on qRT-PCR of SRF targets (*Cald1*, *Acta2*, *Myl9*), luminal progenitor markers (*Kit*, *Cd14*, *Elf5*), *Trps1* and the YAP-SRF joint target *Ctgf* in mammary gland organoids treated for 3 days with Cytochalasin D (2 µM) or DMSO. (o) Schematic of SRF activation in Colony forming assay with sorted luminal progenitor cells. (p) well area covered by colonies formed by luminal progenitor cells treated with cytochalasin D (0.2 µM), Y-27632 (10 µM) or DMSO as control. right side: representative pictures of colonies stained with Giemsa.

When comparing the transcriptome specifically in luminal progenitor cells, we identified 1077 genes (up: 337 genes, down: 740 genes) that were differentially regulated after TRPS1 depletion compared to control (Fig. 5b). The downregulated genes were enriched for processes linked to tissue homeostasis and gland development underlining that TRPS1’s function in the LP population is relevant for the homeostasis of the mammary gland (Fig. 5c). Upregulated genes showed an enrichment for processes related to cytoskeletal dynamics, in particular Actin dynamics (Fig. 5d). In line with that, a TF motif search found Serum Response Factor (SRF)-regulated genes to be specifically overrepresented among the genes that were upregulated after TRPS1 depletion (Fig. 5e). SRF is regulated by actin dynamics and in turn regulates this process through the regulation of cytoskeleton-related genes. This data suggests that TRPS1 is required to restrain SRF activity in LPs. This was further supported by the upregulation of SRF targets in TRPS1-depleted LPs (Fig. 5f, 5h). SRF/MRTF targets were more abundantly expressed in cluster 4 and 0, which contained cells that neither belonged to the LP cluster 2 nor the more mature clusters 5,3,1 (Fig. 5a, 5g). This raises the possibility that TRPS1 is required to repress a transcriptional differentiation program that would normally drive them towards cluster 4,0. We thus next identified marker genes for these clusters exclusively in the shRen samples to identify which genes are strongly expressed in clusters 4,0 under unperturbed conditions (Supplementary table S3). Strikingly, the top marker genes of the more distant cluster 0 contained well-known YAP target genes, such as *Ctgf*, *Cyr61*, and *Klf6*, and cluster 4 genes contained known SRF/MRTF target such as *Actn1* or *Tuba1c* (Fig. 5j). Since YAP/TAZ and SRF/MRTF reciprocally potentiate each other’s activity [22] this implies that TRPS1 fulfills the function to restrain these two pathways which gradually become more active from the LP compartment towards cluster 4 and 0 (Fig. 5i, 5k). CytoTrace analysis on shRen cells revealed that progenitors follow two possible differentiation paths, one leading to mature hormone-responsive ductal cells and a second path leading to non-Hormone-responsive luminal ductal cells which might represent cells that are primed for alveolar differentiation (Fig. 5l).

To investigate how SRF activity affects the mammary gland, we treated mammary gland organoids from WT mice with cytochalasin D (CD) to stimulate SRF activity. After one day of treatment canonical SRF targets were upregulated (Fig. 5m, 5n). Concomitantly, LP marker genes were potently downregulated. Surprisingly, *Trps1* gene expression itself was also downregulated, suggesting that high *Trps1* expression is an inherent feature of the LP gene expression program (Fig. 5n). To test whether this might negatively influence the LP function, we performed a colony forming assay with LP isolated from WT mice and treated them with CD (Fig. 5o). LPs treated with CD completely lost their ability to form colonies showing that elevated SRF activity interferes with LP function (Fig. 5p). In conclusion, these data demonstrate that TRPS1 fulfills an essential function in the LP compartment of the mammary gland by restraining a mechanosensitive YAP/TAZ-SRF/MRTF gene expression program.

### TRPS1 represses Elf5 activity in LP to prevent differentiation

To identify early events that affect the LP population after TRPS1 depletion, we used single-cell ATAC-Sequencing (scATAC-Seq) to identify chromatin regions that become more/less accessible upon TRPS1 depletion. For that, we treated mammary gland organoids from sh*Trps1* mice with doxycycline for seven days to induce the shRNA expression and performed the assay (Fig. S10a). We used matched organoids (Dox- vs. EtOH-treated) to exclude confounding factors due to differences in different organoid preparations. Cells were distributed in eight different clusters (Fig. S10b) and TRPS1-depleted cells showed radically different clustering compared to the EtOH-treated controls (Fig. S10b,c). Based on motif enrichment for lineage-specific transcription factors, clusters 1/2 contained LPs (Elf5), cluster 5 contained basal cells (p63), cluster 7 contained Gata-positive cells, and cluster 8 contained Pgr-positive hormone-sensitive cells (Fig. S10c-g). Most Dox-treated cells were found in clusters 1, 2 and 7 and were specifically excluded from cluster 8. In all clusters, GATA motifs were much more accessible in the Dox-treated condition compared to the control, indicating that the depletion of the GATA-factor TRPS1 leads to a potent opening of its target sites (Fig. S10h). Differential peak analysis showed that TRPS1 depletion indeed had profound effects on the chromatin landscape, as more than 40,000 peaks became more or less accessible (Fig. S10i). The search for motifs which showed most differential accessibility across all differential ATAC-Seq peaks revealed that accessibility to SOX motifs was decreased while ETS1 (family to which Elf5 also belongs) motif accessibility was increased. This indicates that TRPS1 represses ETS TF activity and stimulates SOX TF activity. Sox genes mainly Sox4 but also Sox5 and 12 have been shown to promote BC development and progression [23].

These data reveal TRPS1 as a crucial regulator of chromatin accessibility in mammary gland cells, and they suggest that one function of it is to keep Elf5 activity low. This would be consistent with TRPS1’s role in LP function since unrestrained Elf5 activity is known to drive luminal progenitors into alveolar cell maturation [24].

### TRPS1 represses SRF indirectly via the repression of RhoA modulators

To investigate how TRPS1 maintains the LP program at the target gene level, we identified direct TRPS1 targets in sorted luminal cells using CUT&RUN. 725 high confidence peaks were identified among which ∼70% were in promoters (Fig. 6a,b and S11). Interestingly, the genomic regions bound by TRPS1 showed enrichment for C2H2 zinc finger motifs (Fig. 6c) suggesting that in luminal cells, gene regulation by TRPS1 relies more on its C2H2 zinc fingers than its GATA binding domain. This contrasts with TRPS1’s binding pattern in breast cancer cells where i) TRPS1 almost exclusively binds at enhancers, and ii) TRPS1 peaks are strongly enriched for GATA motifs but not C2H2 zinc finger motifs [3, 11].

**Figure 6:**
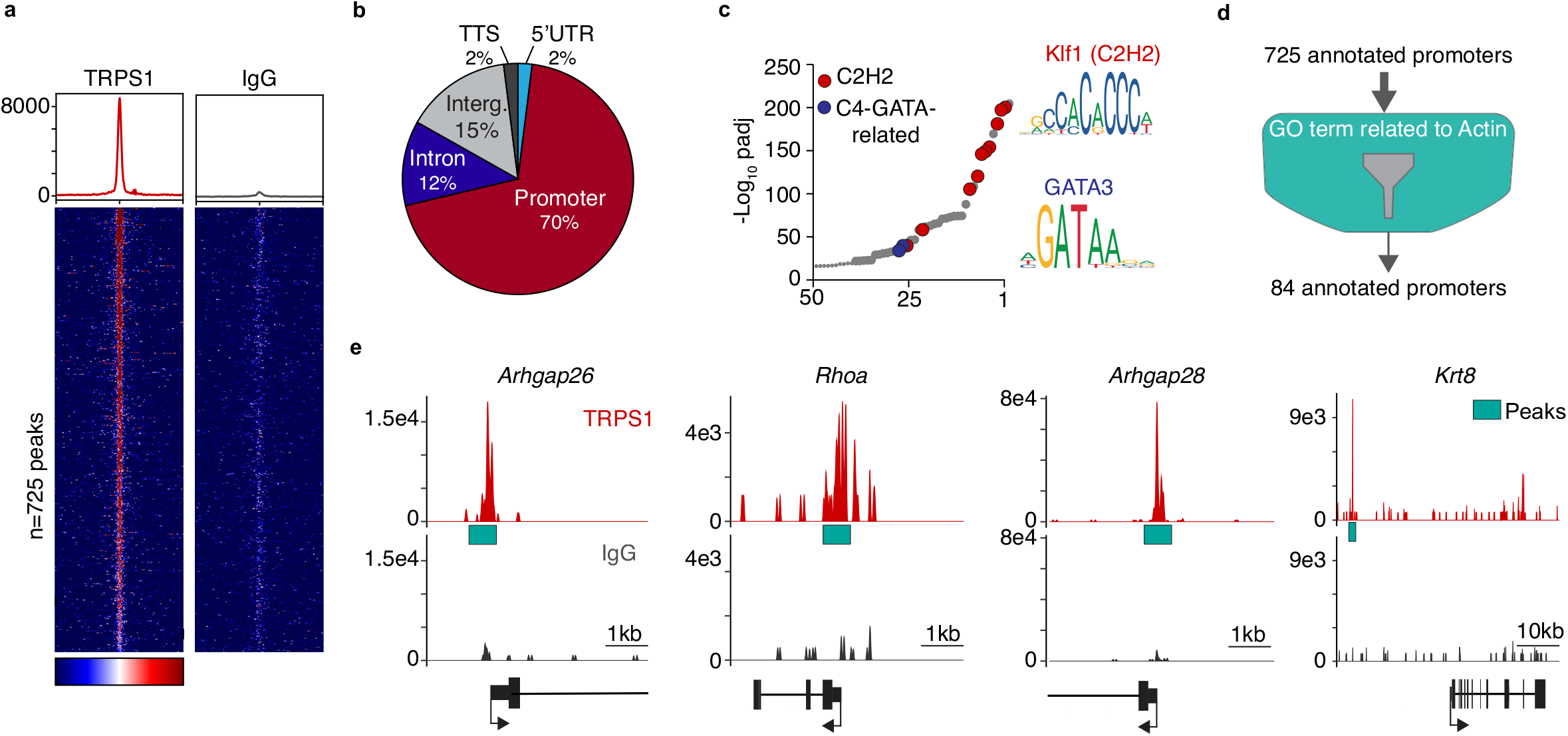
TRPS1 represses SRF/MRTF activity via indirect modulation of RhoA activity. (a) Heatmap of TRPS1 Cut&Run binding peaks in promoters (n=725). IgG serves as negative control. (b) Venn diagram of the distribution of TRPS1 binding peaks in the genome. TSS = Transcription Start Site; Interg. = Intergenic region; UTR = Untranslated region. (c) Scatter plot for the enrichment of C2H2 and GATA binding motifs in regions bound by TRPS1 identified by Cut&Run. (d) Schematic of the strategy used to select TRPS1-target genes involved in the regulation of actin dynamics. (e) Cut&Run genomic tracks for TRPS1 binding (Red) and control IgG (Gray) centered on the promoter regions of the indicated genes. Green boxes mark the position of the Cut&Run peaks.

We then filtered the corresponding gene promoters based on their GO term annotation and found 84 genes whose function was linked to “Actin cytoskeleton” dynamics (Fig 6d and S4). A large subset of these genes encoded modulators of RhoA activity or proteins whose function is regulated by RhoA including *Rhoa* itself*, Arhgap26*, *Arhgap28*, *Krt8* (Fig 6d,e) *Atf4*, *Tip1*, *mTor* [25–29]. This set of genes might be targeted by TRPS1 to indirectly influence RhoA activity. Since RhoA functions upstream of SRF/MRTF, this would result in the repression of SRF/MRTF activity (Figure 7).

**Figure 7:**
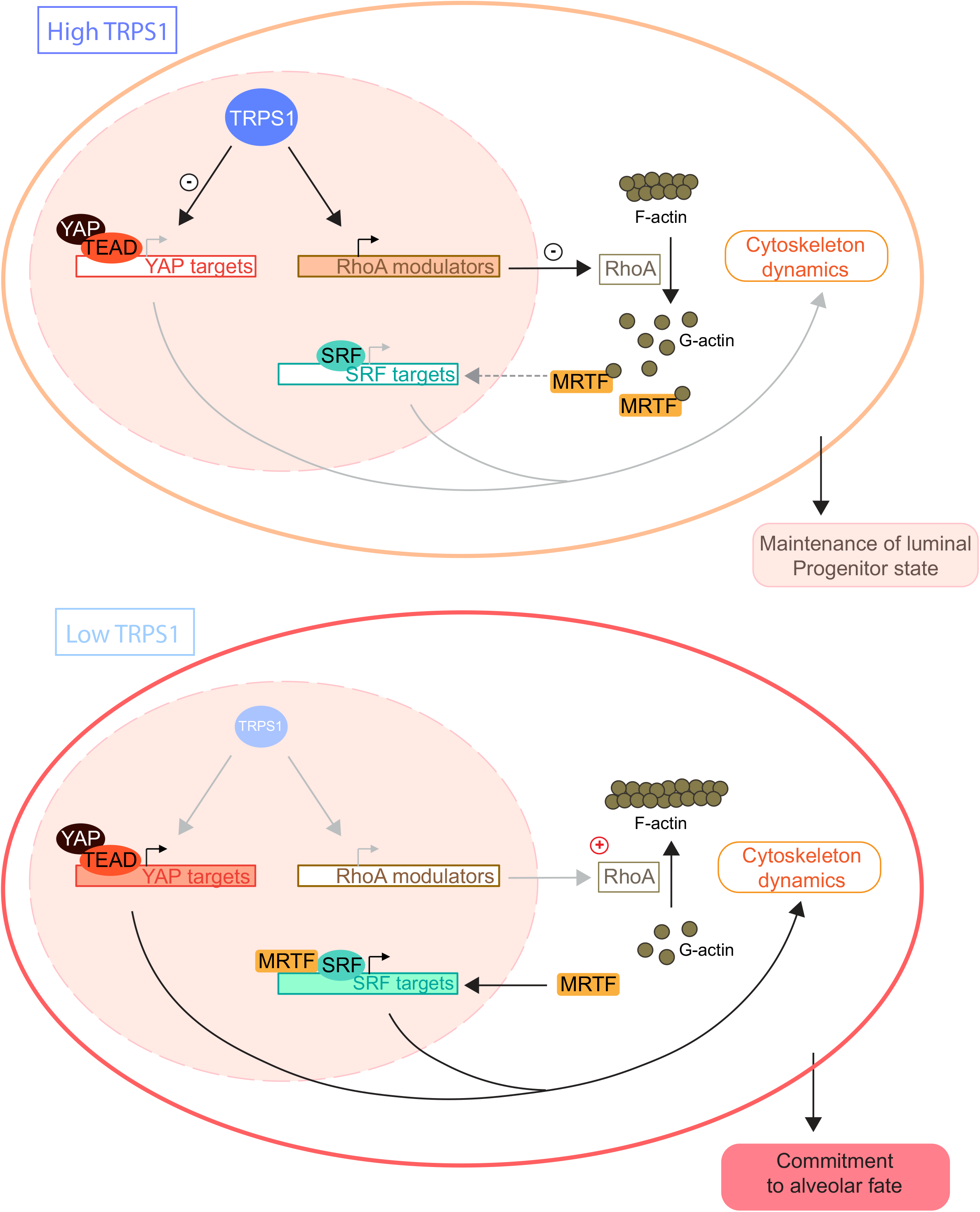
Model for how TRPS1 prevents luminal progenitors from entering the alveolar differentiation path. In luminal progenitor cells, TRPS1 activity is high and represses both YAP and SRF/MRTF activity: YAP repression results from the direct modulation of YAP target genes while SRF/MRTF activity is repressed indirectly via the modulation of RhoA activity. In luminal cells committed to the alveolar fate, TRPS1 activity is low, the repression over SRF/MRTF and YAP activities is lifted and forces the cells to exit the progenitor state.

## Discussion

In this work we have developed a mouse model allowing the conditional ubiquitous depletion of TRPS1 using dox-inducible shRNA expression. We found that this model provided a way to ubiquitously diminish TRPS1 function without affecting the skeleton and cartilage development and early mammary gland development since it allows TRPS1 depletion in the adult animals. TRPS1 has recently emerged as an oncogene in specific breast cancer types. This work thus provides the first indication that targeting TRPS1 with a drug would not result in serious side effects. Indeed the detailed phenotypic analysis did not reveal any major health issues upon TRPS1 depletion in the female mice who remained healthy over ten weeks of doxycycline treatment. In the future, it will be interesting to see how shTRPS1 mice respond in a breast cancer mouse model. Here, however we focused on the physiologic functions of TRPS1, mainly in the mammary gland.

Consistent with TRPS1 expression being restricted to luminal cells, basal cells barely responded to TRPS1 depletion, whereas the transcriptome of luminal cells-particularly the luminal progenitor compartment-was markedly affected. Consistently, TRPS1 depletion had a negative impact on LP function, e.g. in CFC assays, demonstrating that TRPS1 is essential for these cells. Unexpectedly, we observed that GFP-negative luminal cells from shTrps1 animals formed a higher number of colonies than shRen controls. This suggests that compensatory mechanisms are present *in vivo* to overcome the reduced functionality of TRPS1-depleted LPs. This could explain why we found no apparent changes of the mammary gland histological structure after long-term TRPS1 depletion.

Searching for regulatory pathways that are modulated by TRPS1 to maintain the LP pool and their functionality, we found that TRPS1 represses the SRF/MRTF regulatory pathway linking actin dynamics to SRF-mediated gene transcription. Notably, low SRF activity is crucial to maintain the LP function since hyperactivation of SRF in luminal cells did not only lead to repression of LP marker genes but also completely abolished colony forming abilities in CFC assays whereas ROCK inhibition promoted colony formation.

Our scRNA-Seq pseudotime analysis revealed that the LPs can follow two possible paths of differentiation one leading to mature hormone-sensing cells and the other to a population that could represent hormone receptor-negative committed alveolar precursors. The alveolar branch shows gradual activation of SRF/MRTF as well as YAP/TAZ suggesting that there might be already a priming mechanism in alveolar precursors that would require SRF and YAP activity. In virgin mice, a small number of alveolar cells are differentiated at each estrus cycle, and this number increases during pregnancy [30]. Although in the virgin gland YAP is mainly active in basal cells, at pregnancy it becomes very active in proliferating alveolar cells prior to differentiation of the alveoli [31]. MRTF is crucial to maintain basal cell contractile function during lactation [32], but there is no report of SRF/MRTF playing a role in alveolar differentiation of the luminal cells. However, SRF and YAP are both involved in mechano-transduction and they have overlapping targets and potentiates each other’s activity [22] so they could cooperate to induce differentiation of alveolar cells. The development of Alveoli and the associated changes in cell morphology is a process that is likely to produce mechanical strain and therefore influence the actin cytoskeleton/SRF activity not only in the basal layer but also in the adjacent luminal alveolar cells [33]. YAP/TAZ activity in luminal cells could also be influenced by mechanical strain, as it is the primary signal stimulating YAP/TAZ activity in the liver [34]. Keeping YAP/TAZ activity in check is crucial in luminal cells since uncontrolled YAP has been shown to trigger overgrowth of aberrant luminal cells expressing basal markers that lead to altered ductal morphology and basal-like BC phenotypes [35]. In any case, TRPS1 needs to repress YAP/TAZ and SRF/MRTF activity to preserve the LP pool by preventing LPs from entering the alveolar differentiation path. Interestingly, YAP/TAZ is repressed directly by TRPS1 by binding to joint sites whereas SRF/MRTF repression seems to be largely indirect, e.g. via repression of a set of genes involved in modulating RhoA activity so that independent mechanisms can assure their stable repression.

In support of the hypothesis that TRPS1 prevents differentiation, our scATAC-Seq experiments to dissect the early events following TRPS1 depletion in mammary gland organoids revealed a strong activation of Elf5. Elf5 overexpression has been shown to support the differentiation of luminal progenitors into alveolar cells [24]. Concomitantly, GATA motifs become more accessible and this could result in the activation of GATA targets as these sites could be more accessible to GATA activators like GATA3 which is also essential for the differentiation of luminal progenitor cells along the secretory alveolar sublineage [19, 20]. Taken together our results suggest that TRPS1 maintain the LP population mainly by preventing entry in the alveolar differentiation process. This involves the repression of a YAP/SRF mechano-transduction program (Figure 8).

In our CUT&RUN experiments from freshly sorted mouse mammary gland cells, we observed a strikingly different binding behavior of TRPS1 compared to ChIP-Seq experiments from cultured breast cancer cells [2, 3, 36]. Whereas in breast cancer cells, TRPS1 mainly bound to enhancer regions via its GATA domain, our CUT&RUN demonstrate a promoter-proximal binding pattern that is potentially mediated by TRPS1’s C2H2 zinc fingers. Whether this is due to technical differences of ChIP-Seq vs. CUT&RUN or whether TRPS1 binding is altered during malignant transformation needs to be addressed in the future. This data imply, however, an important function of TRPS1’s C2H2 zinc fingers for mammary gland function and could potentially explain why no phenotype has been observed in TRPS1 patients where only the GATA zinc finger region is affected.

In young BRCA1/2 carriers [37] and aged women [13, 38], there is an expansion of aberrant LP population expressing basal traits. These are believed to be the cells of origin of BC and therefore their expansion highly increases the risk to develop breast cancer. Since TRPS1 maintains a functional LP population in the homeostatic mammary gland, it is tempting to speculate that TRPS1 helps to drive luminal progenitor expansion. If so, targeting TRPS1 already at an early stage could be an effective preventive and safe strategy for BRCA1/2 carriers – as evidenced by the ubiquitous shTRPS1 mouse model.

In summary our data demonstrate a crucial role of TRPS1 in LP function of the mammary gland, and we have evidence that targeting TRPS1 could be a promising target for future therapy.

## Materials and methods

A list of reagents (antibodies, chemicals, cytokines, commercial kits) is provided as Supplementary Table 5. For a detailed description of the experimental procedures including the phenotypic analyses carried out at the german mouse clinic, western blotting, Immunofluoresence, qRT-PCR, RNA-Seq and CUT and RUN See SI Appendix

### Mice

CAG-lsl-RIK, TRE-tGFP-shTrps1#1, shTrps1#2 or shRenilla [39] (Premsrirut, 2011), and CAG-rtTA3 mice lines were obtained from the Mirimus Company (NY, USA). The shRNA sequences can be found in Supplementary Table 6. Details about the husbandry and the breeding of experimental animals can be found in the SI Appendix and Figure S1. Mice were used for experiments within an age range of 12 to 20 weeks. To induce shRNA expression, the animals were fed with doxycycline-supplemented food (ssniff-Spezialdiäten company, Germany).

### Sorting mammary epithelial cells by flow cytometry

The mammary gland epithelial cell isolation method is described in detail in SI Appendix. Mammary gland cells were incubated with a mixture of biotin-conjugated lineage antibodies (anti-TER-119, anti-CD31 and anti-CD45, see Table S5). After washing in PBS 2% FBS, the cells were stained with fluorophore conjugated antibodies (PE-Anti-mouse EpCAM, APCFire750-anti-mouse CD49f and APC-anti - ckit, PEvio770-anti mouse CD14) and strepatavidin-eFluor450 (Table S5). For flow cytometric analysis and sorting, stained cells were washed twice, resuspended in PBS 2% FBS, filtered (40 µm cell strainer) and 1 μM SYTOX Blue dead cell stain (Thermo Fisher Scientific) was added. For further details about the sorting procedures see SI Appendix and Supplementary Figures S4-S9 and S11.

### CITE-Seq

Mammary gland cells were isolated as described in SI Appendix section CITE-Seq. 0.5 ug of each Total Seq antibodies (anti-EpCAM and anti-CD49f, Table S5) was added to the cell suspension and the samples were incubated 30 min at 4°C. Cells were washed in PBS + BSA 2% + Tween 20 0,01%, counted and adjusted to a concentration of 1000 cells/ µL. Cells were subjected to the Chromium single cell 3’ assay v3 (10X Genomics) as recommended by the manufacturer. After cDNA amplification, ADT-derived cDNAs and mRNA-derived cDNAs were separated based on their size using 0.6x AMPure XP Beads (Beckman Coulter). The mRNA-derived cDNAs contained in the bead fraction were further processed following the standard 10X Genomics protocol in order to generate single-cell (sc)RNA libraries. The ADT-derived cDNAs contained in the supernatant were further purified (see SI Appendix) and used as template in a PCR reaction with the NEBNext^®^ High Fidelity Master Mix (NEB), a Truseq small RNA RPIx (containing i7 index) primer and the 10X Genomics SI-PCR primer (see Table S6) to generate the ADT sequencing library. The scRNA-Seq libraries and the ADT libraries were then sequenced together on an Illumina NextSeq500 platform (details in SI Appendix).

#### Mammary gland organoids isolation and cultivation

Freshly isolated mouse mammary gland tissue was finely chopped and digested in 2.5 mL of Leibovitz’s L-15 Medium (Thermo Fisher Scientific) supplemented with 3 mg/mL Collagenase A and 1.5 mg/mL Trypsin in a GentleMACS^TM^ Octo dissociator (Miltenyi) for 1h at 37°C, 100 rpm. After centrifugation, the cell pellet was resuspended in DMEM/F12 medium, filtered (70 µm cell strainer) and centrifuged again. Cells were resuspended in Matrigel (Corning) dispensed in 10 µL-drops in a 48-well plate and cultured in Organoid medium (DMEM/F12 + Glutamax with addition of 1X N-2 Supplement, 1X B27 Supplement, 100 ng/mL Neuregulin, 100 ng/mL Noggin and 100 ng/mL R-spondin) at 37°C, 5% CO_2_. Organoids were passaged every 2 to 3 weeks. Details about passaging and dissociating organoids are provided in SI Appendix.

### Material for qRT-PCR and RNA-Seq

For qRT-PCR, 3,000 sorted luminal progenitor/mature cells and about 50,000 isolated mammary organoid cells were used per sample for RNA extractions (Fig S6). RNA from Luminal mature and progenitor cells was amplified using the “NEBNext^®^ Single Cell/Low Input cDNA Synthesis & Amplification” (NEB). For RNA-Seq 20,0000 to 50,000 sorted basal and luminal cells from the shTrps1 ubiquitous mouse model (Fig S4) and about 1,000 to 15,000 sorted GFP^+^ mKate2^+^ sorted luminal cells from the ShTrps1 luminal-specific mouse model (Fig S5) were used for RNA isolation and subsequent library preparation. For CUT and RUN, 200,000 sorted luminal cells from wild type mouse were usd per reaction.

### Colony forming assay

Luminal or luminal progenitor cells were isolated from mouse mammary glands and sorted in 96-well plate filled with 5% FBS EpiCULT full medium (STEMCELL Technologies). 500 luminal cells or 200 luminal progenitors were then transferred to a 48-well plate previously seeded with 10,000 NIH3T3 irradiated feeder cells per well and incubated at 37°C, 5% CO_2_. 24h later the medium was exchanged for EpiCULT full medium without FBS. When needed, doxycycline (10 µg/mL), ethanol (1:1000), cytochalasin D (0.2 μM) or the ROCK inhibitor Y-27632 (10 μM) were added to the medium. Cells were incubated for 7 days and medium was exchanged every 2-3 days. Colonies that ultimately formed were fixed for 1 min in methanol and stained in 10% Giemsa staining solution for 20 min.

### scATAC-Seq

Nuclei suspensions from 60,000 cells isolated from mammary gland organoids were processed using the 10x Genomics Chromium Controller and Chromium Next GEM Single Cell ATAC Reagent Kits v2 according to the manufacturer’s protocol (CG000496). 7600 nuclei were loaded onto the Chroimium Controller to recover 5000 nuclei for library prepration and sequencing. Total yield and quality of cDNA was assesed on DNA 7500 assay (Agilent 2100 Bioanalyzer). The libraries were pooled and sequenced on Illumina NovaSeq 6000 System in combination with SP 100 cycles v1.5 kit. The following sequencing cycles were performed: R1 (sequencing of interest: 51 bp; R2 (sequencing of interest): 51 bp; i7 index (sample index): 8 bp; i5 index (10X barcode): 16 bp. Extraction of FastQ files was done using bcl2fastq v2.20.0.422 (Illumina). More details about procedure and data analysis is provided in SI Appendix.

## Data Availability

Source data are provided with this paper upon request. The Next-generation sequencing data generated in this study have been deposited in the GEO database under accession code GSE243511.

## Acknowledgements

B.v.E. was supported by grants from the BMBF (16GW0271K), DFG (EY 120/4-1), and the German Cancer Aid (Deutsche Krebshilfe; 70113138). The FLI is a member of the Leibniz Association and is financially supported by the Federal Government of Germany and the State of Thuringia. The DNA Sequencing, the Proteomics, and the Flow Cytometry core facilities as well as the core service histology of the FLI are gratefully acknowledged. We would like to thank all the members of the von Eyss lab, von Maltzahn lab, and Kaether lab for helpful discussion and Christin Ritter and Tom Hünniger and all the animal care takers at FLI for excellent technical support. Some of the figures were created with BioRender.com

---

**Supplementary Figure S1:**
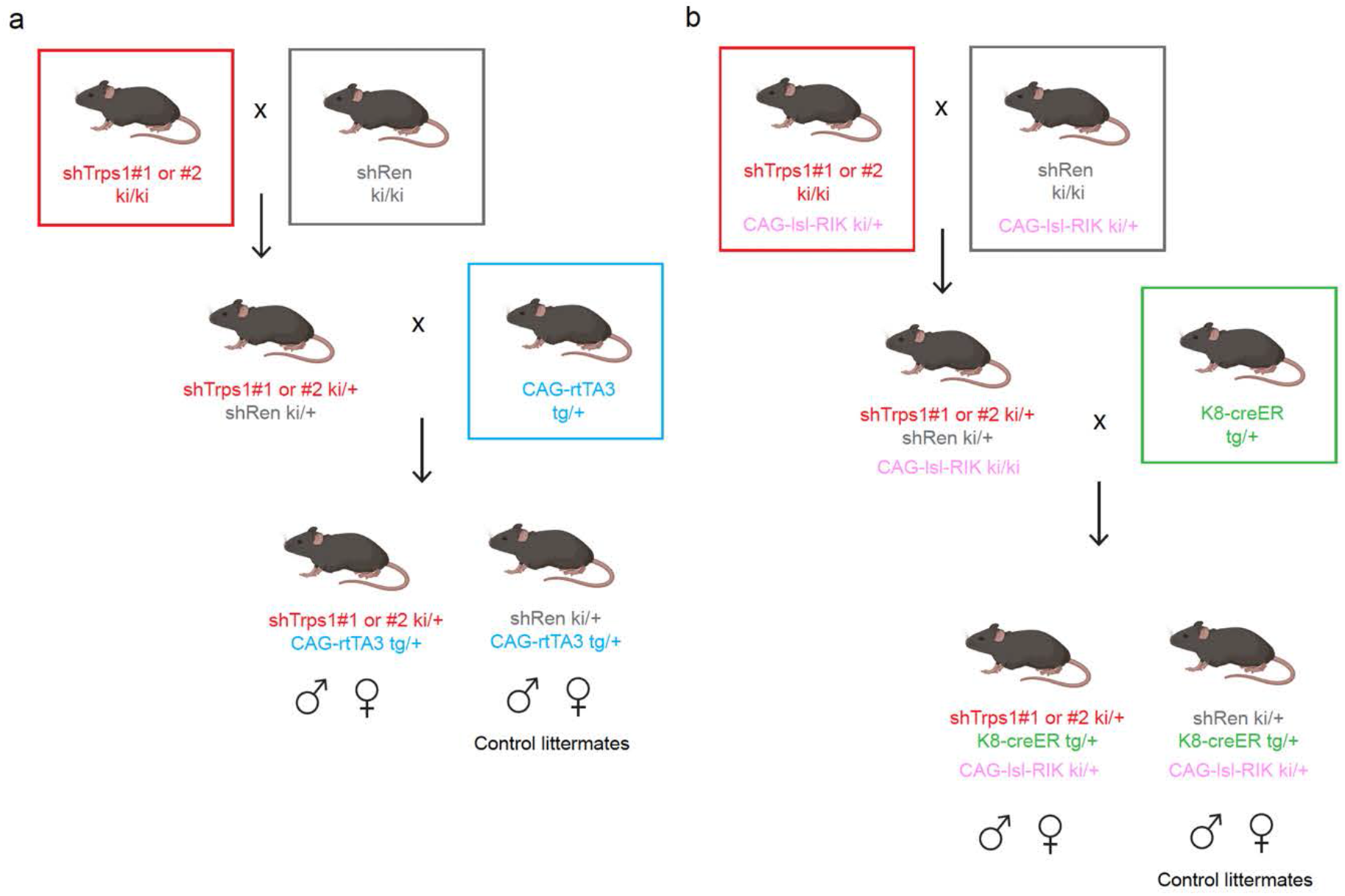
Breeding Schemes. (a) generation of animals ubiquitously expressing shTrps1: homozygous shTrps1 (ki/ki) are crossed with homozygous shRen (ki/ki) to obtain progeny carrying both shTrps1 (ki/+) and shRen (ki/+) alleles. These animals are then crossed with CAG-rtTA3 heterologous (tg/+) animals to obtain shTrps1 ki/+, CAG-rtTA3 tg/+ experimental animals and shRen ki/+ CAG-rtTA3 tg/+ control littermates. generation of animals expressing shTrps1 specifically in the luminal compartment: animals homozygous for shTrps1 (ki/ki) and carrying a CAG-lsl-RIK allele (ki/+) are crossed with animals homozygous for shRen (ki/ki) and carrying a CAG-lsl-RIK allele (ki/+) to obtain progeny carrying both shTrps1 (ki/+) and shRen (ki/+) alleles and homozygous for CAG-lsl-RIK (ki/ki). These animals are then crossed with K8-CreER (tg/+) animals to obtain shTrps1 ki/+, CAG-lsl-RIK ki/+, K8-CreER tg/+ experimental animals and shRen ki/+, CAG-rtTA3 tg/+, K8-CreER tg/+ control animals.

**Supplementary Figure S2:**
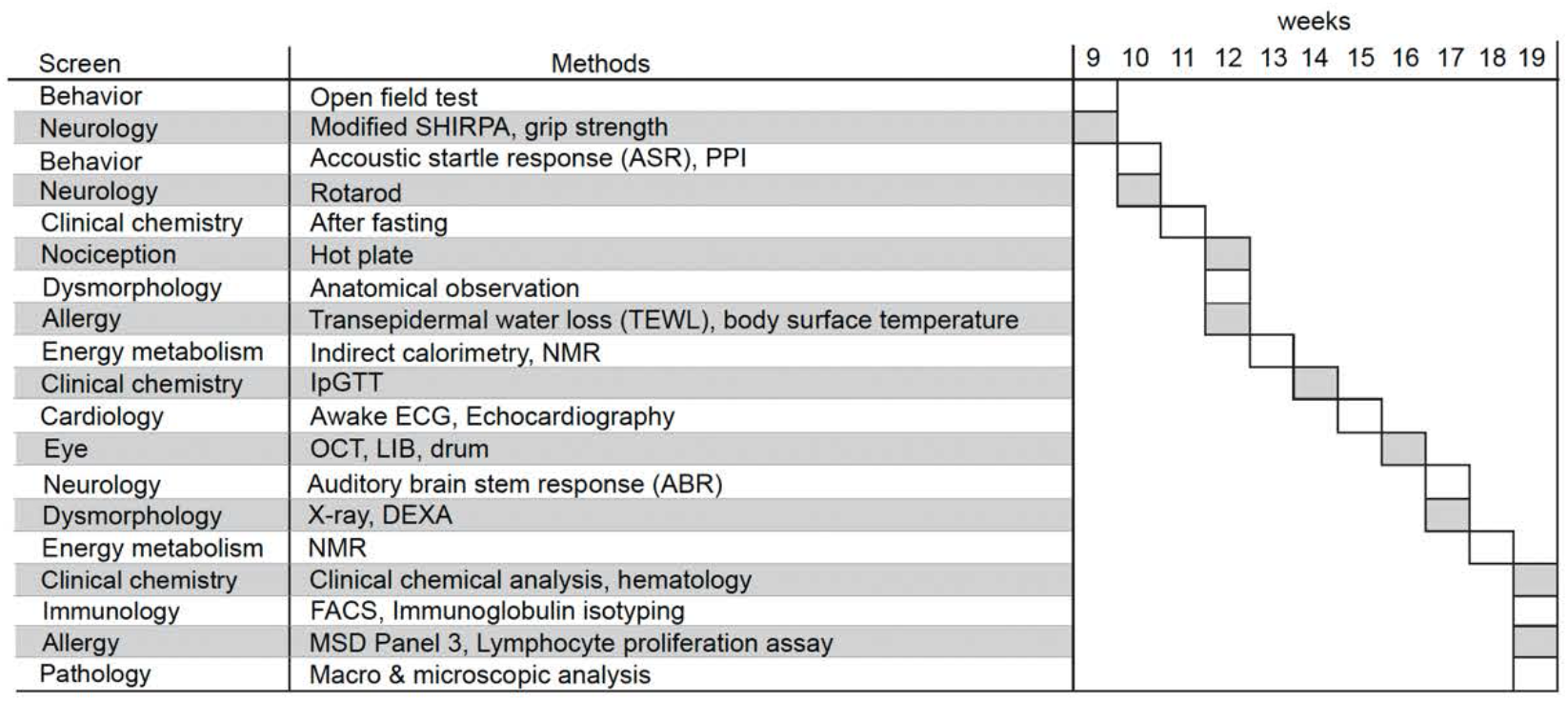
Mouse Phenotypic analysis timeline.

**Supplementary Figure S3:**
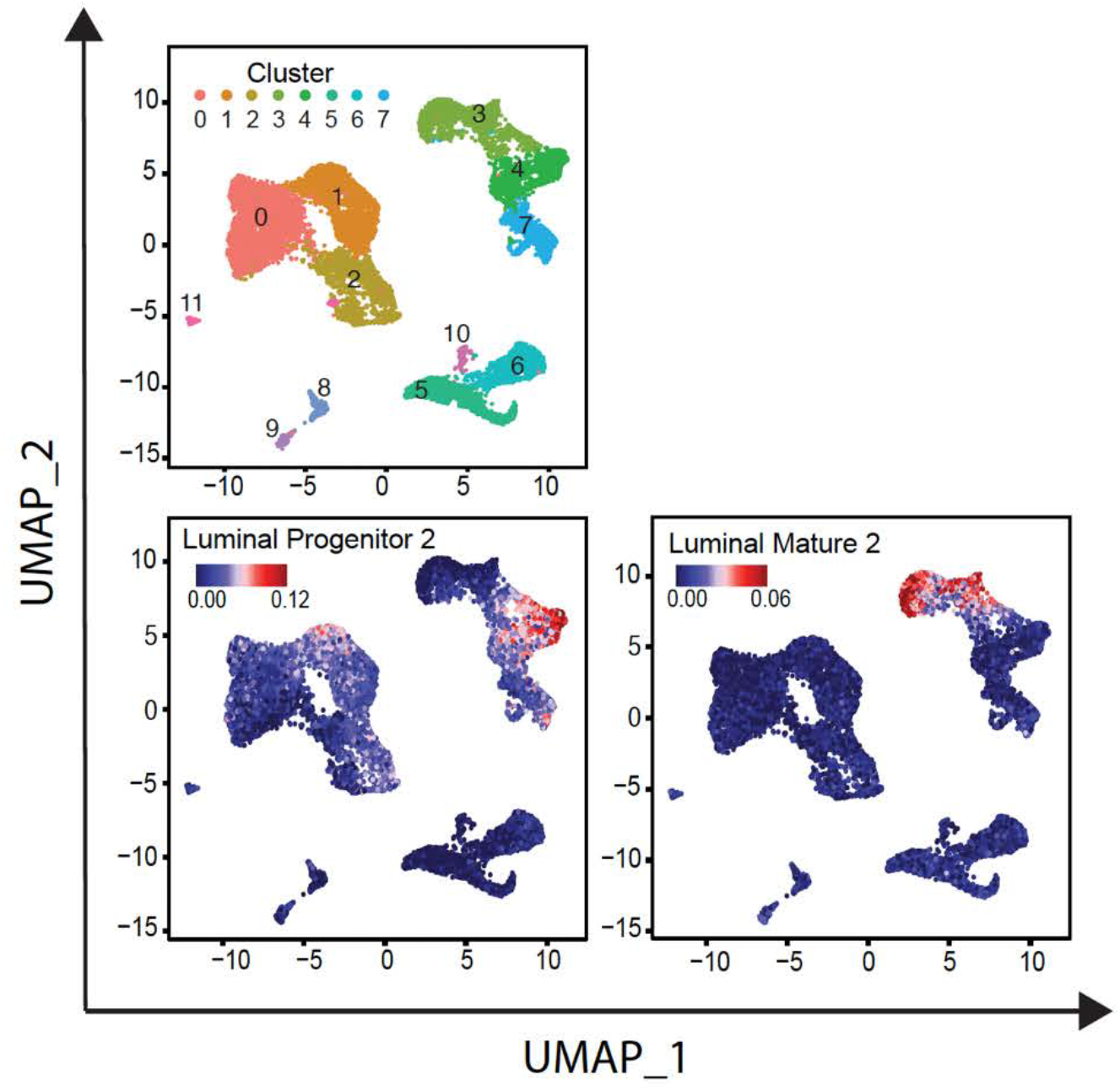
UMAP dimensionality reduction plots of the CITE-Seq data from shTrps1 and shRen mammary gland cells combined (see Figure 3) showing the gene set activity based on AUCs of a luminal progenitor and a luminal mature gene set published by Lim et al. (2010).

**Supplementary Figure S4:**
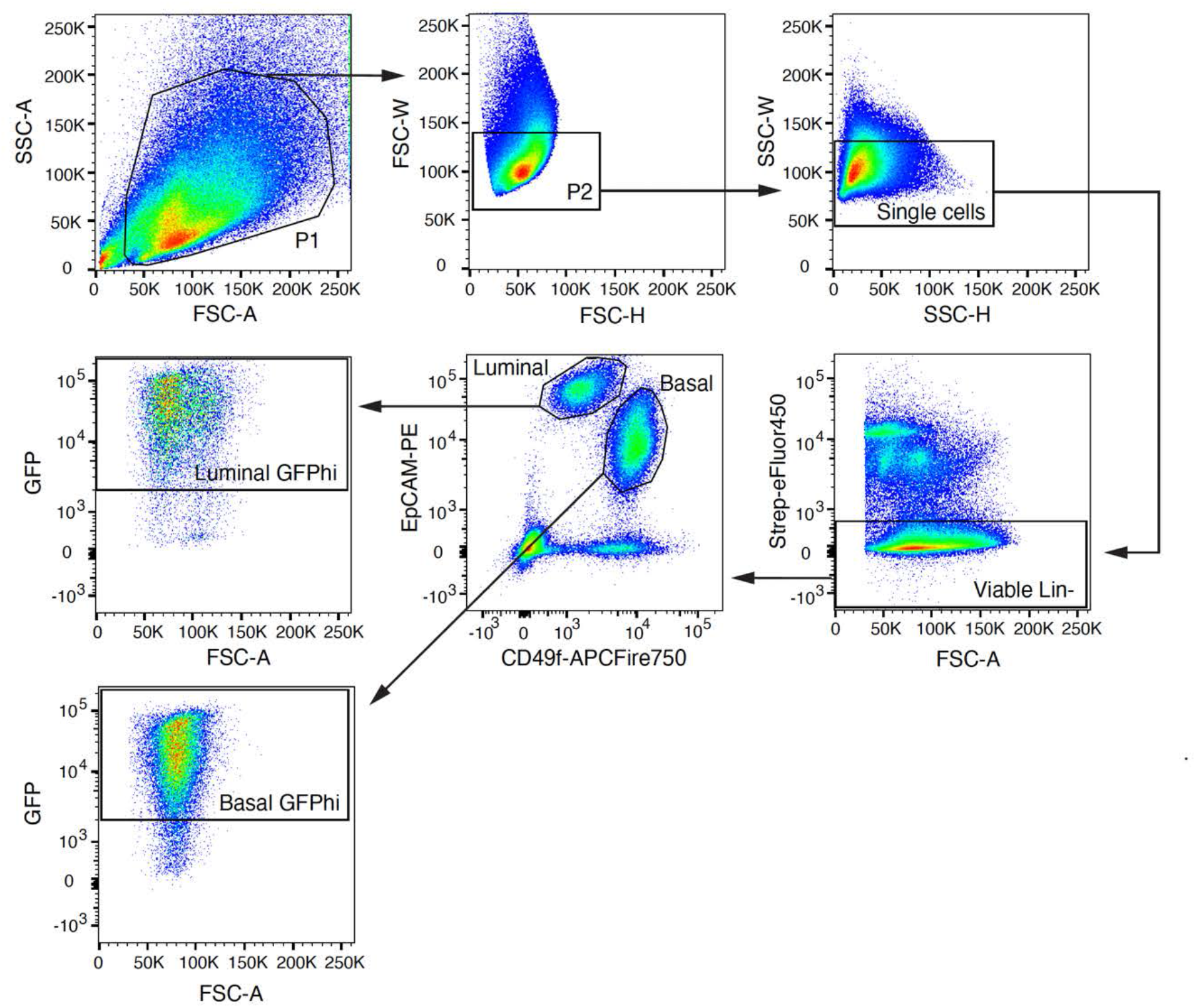
sorting strategy used in Figure 2c to isolate basal and luminal cells expressing shTrps1 (GFPhi) from ubiquitous shTrps1 model mice.

**Supplementary Figure S5:**
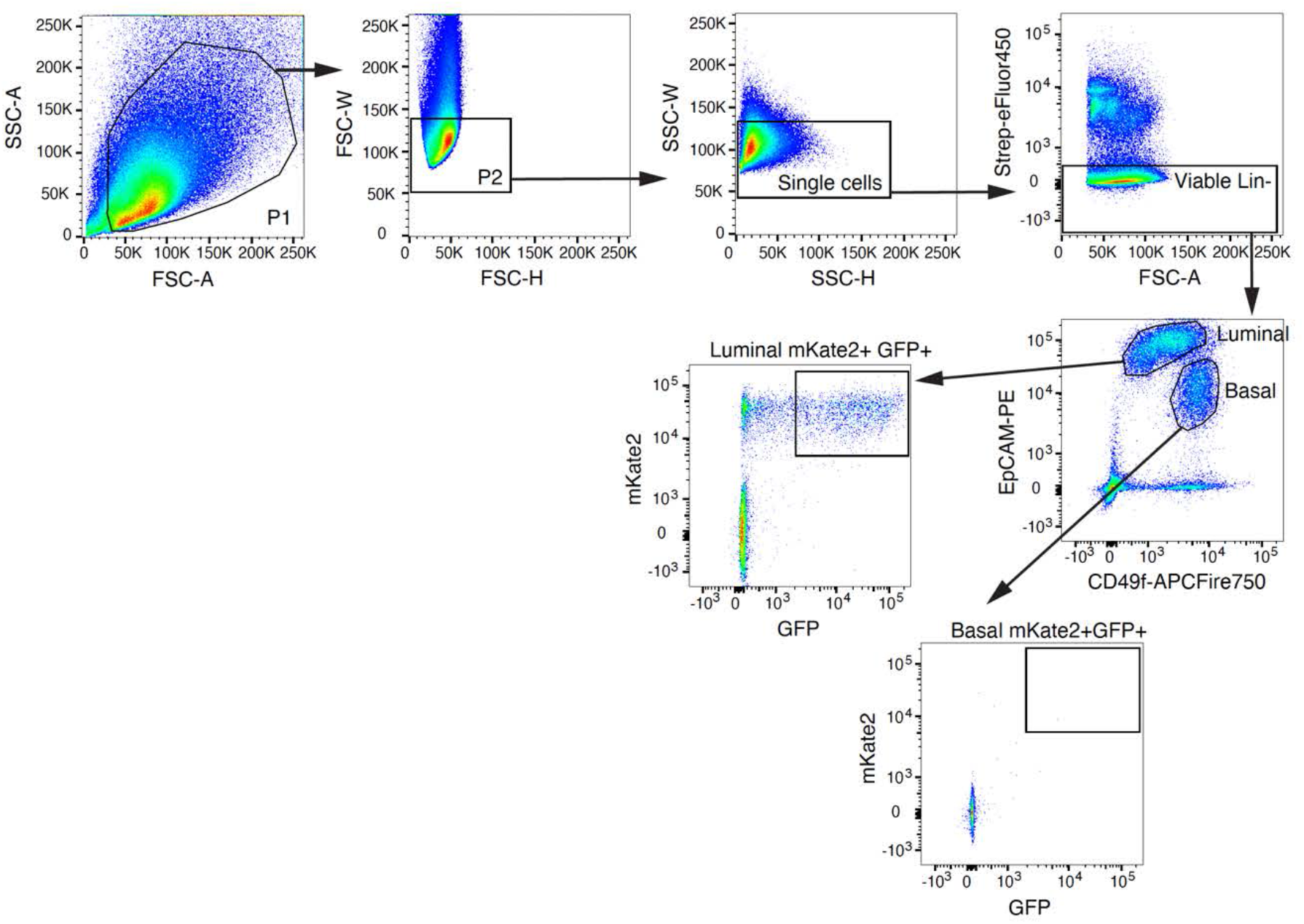
sorting strategy used in Figure 2l to isolate mammary basal and luminal cells expressing shTrps1 (mKate2+ GFP+) from luminal-specific shTrps1 model mice.

**Supplementary Figure S6:**
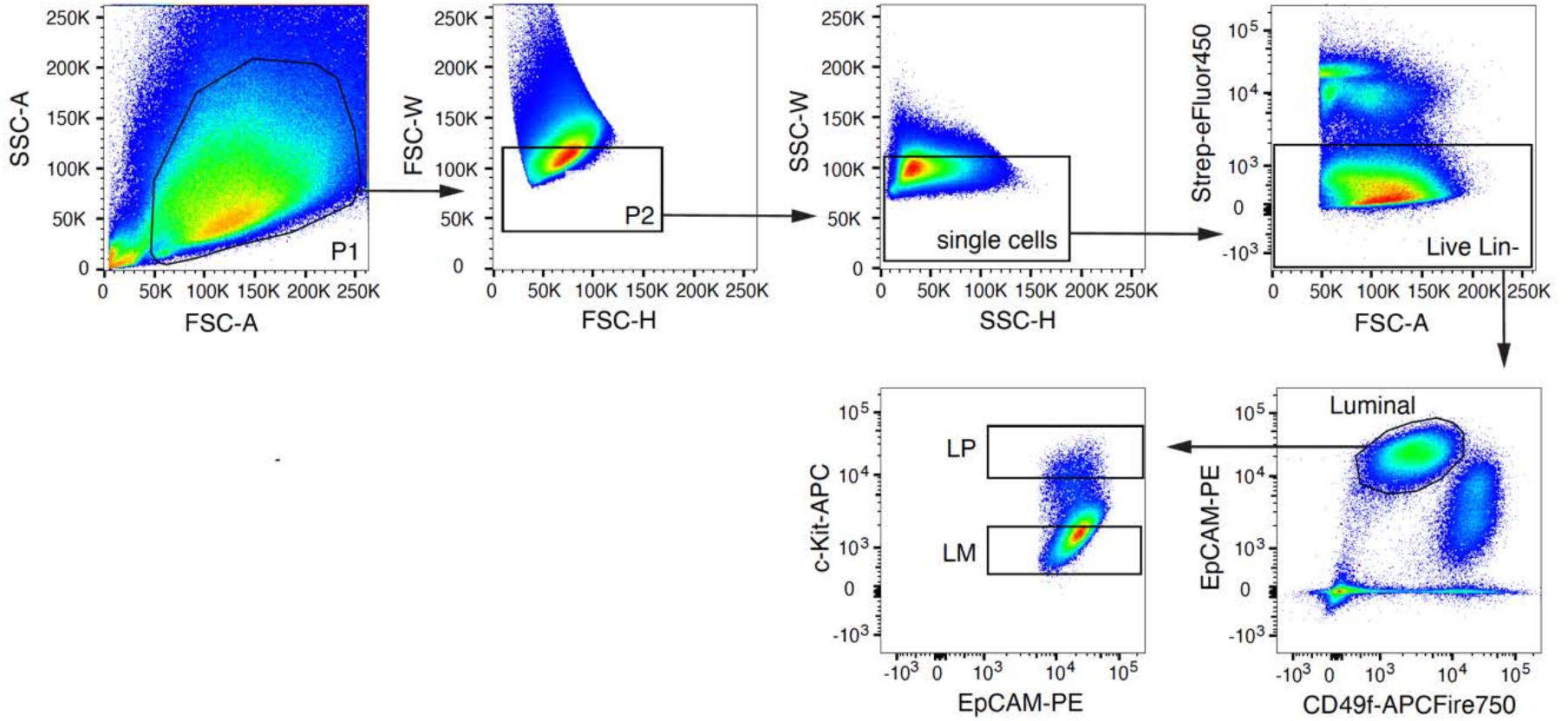
sorting strategy used in Figure 3 to isolate mammary progenitor and mature luminal cells from wild type mice.

**Supplementary Figure S7:**
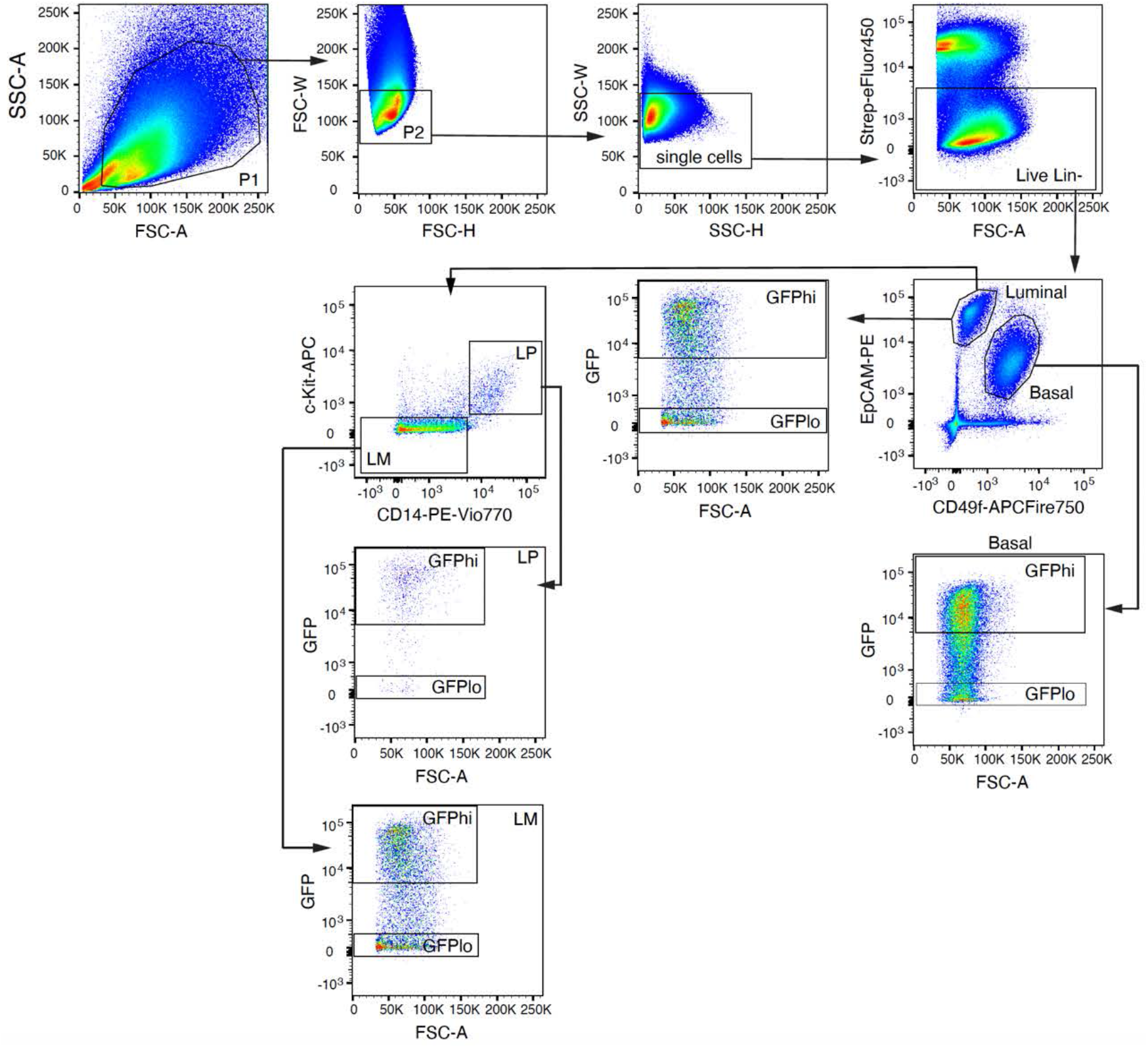
gating strategy used in Figure 4a to analyze the proportion of GFPhi and GFPlo cells in the different mammary epithelial subpopulations: Basal, Luminal, Luminal mature (LM) and Luminal progenitor (LP).

**Supplementary Figure S8:**
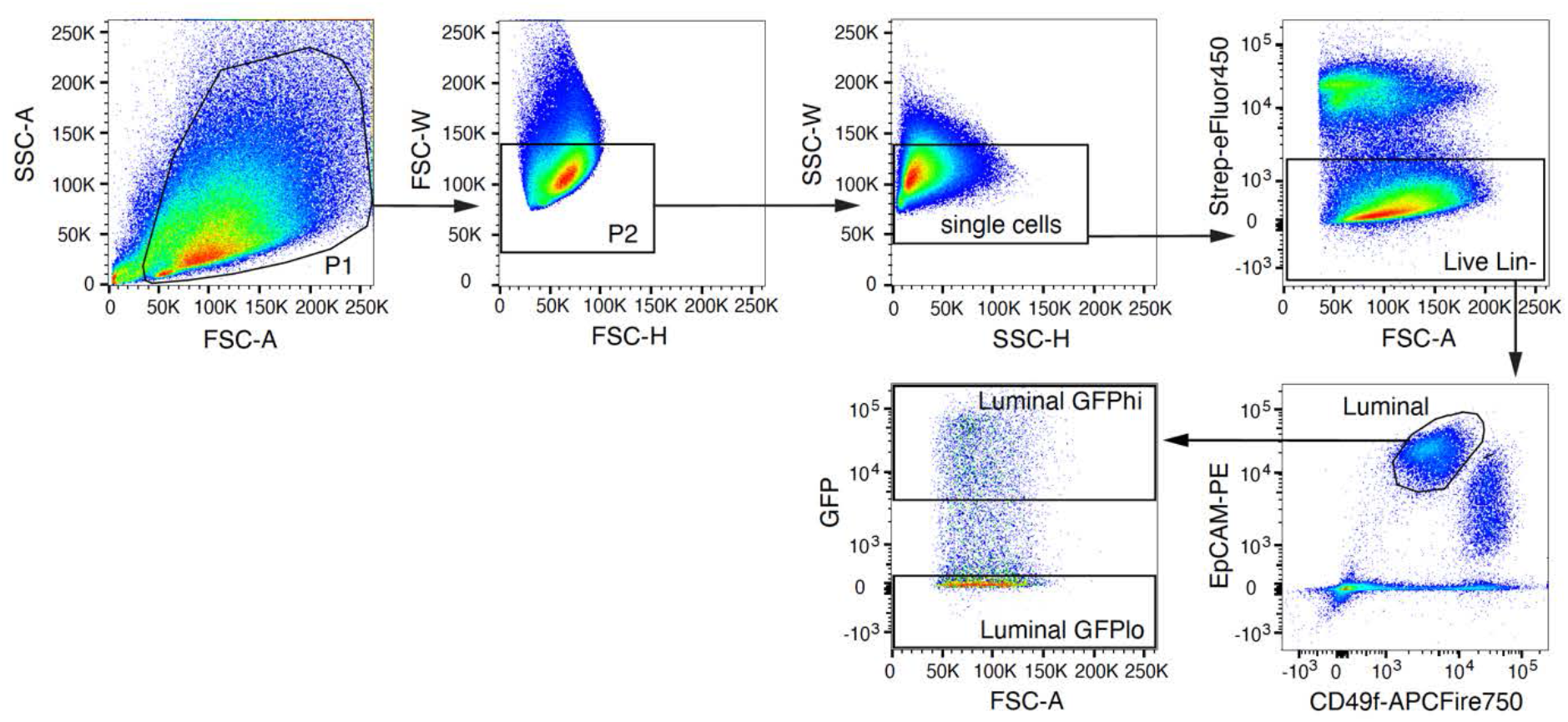
sorting strategy used in Figure 4d-e to isolate mammary luminal cells expressing high shTrps1 (GFPhi) or expressing low of no shTrps1 (GFPlo) for a colony forming assay.

**Supplementary Figure S9:**
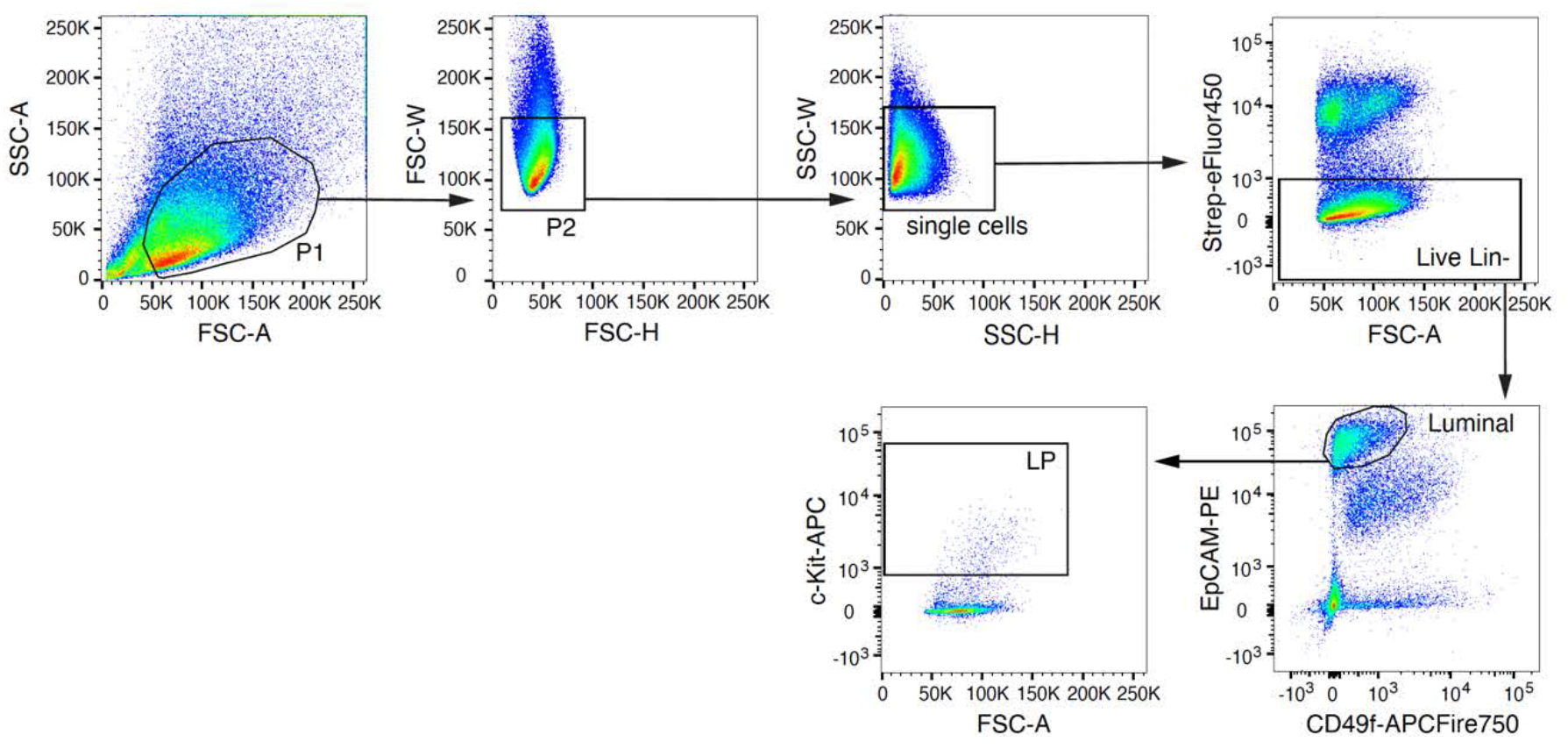
sorting strategy used in Figure 5o-p to isolate mammary luminal progenitor cells from wild type mice for colony forming assay.

**Figure S10:**
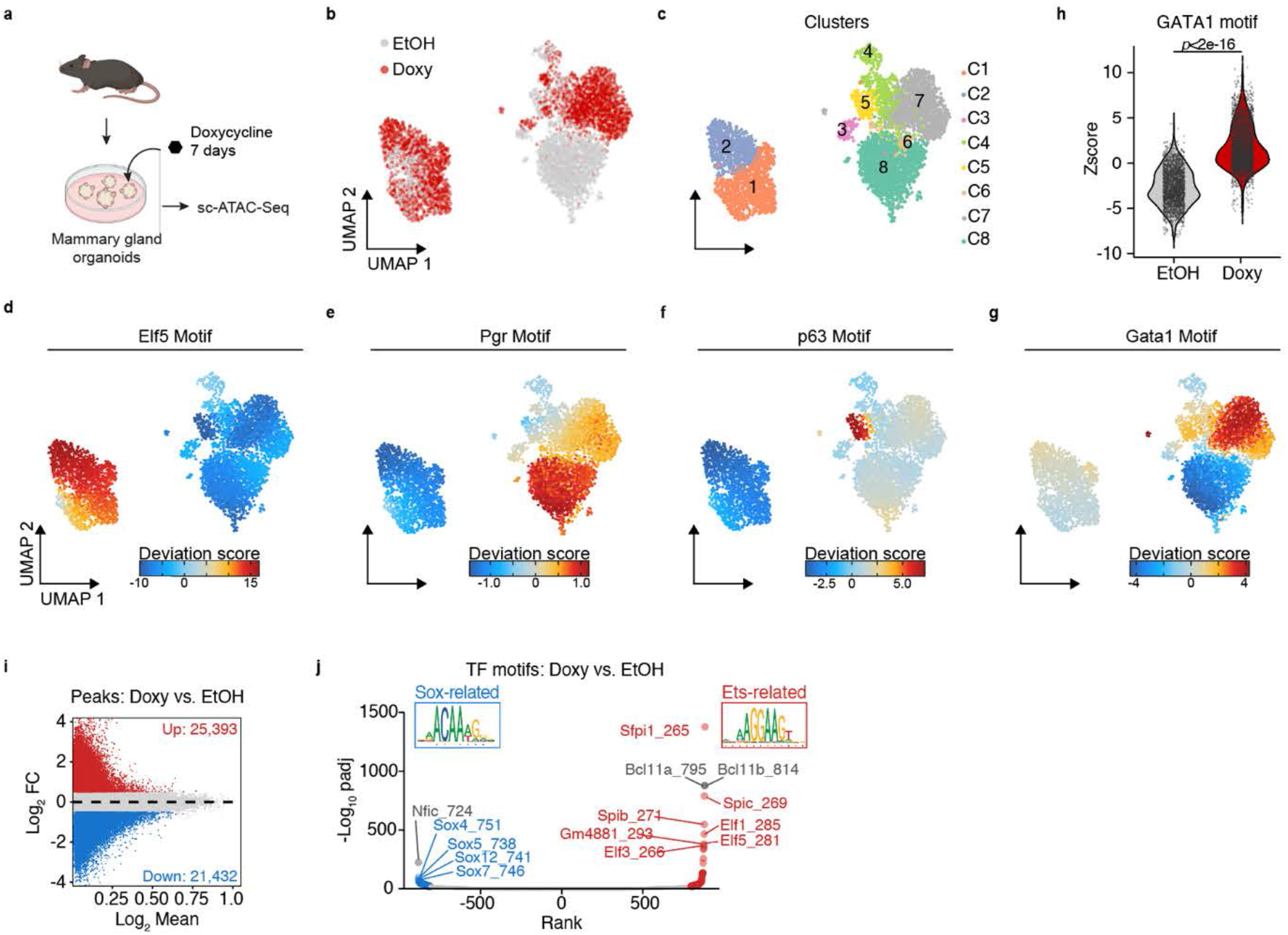
TRPS1 prevents LP differentiation. (a) Experimental design: organoids prepared from mammary glands isolated from wt BL6 mice were treated with doxycycline for 7 days and subjected to sc-ATAC-Seq (b)-(c) UMAP dimensionality reduction plots of the mammary organoid cells. (b) Cells treated with Dox or Ethanol are highlighted in red and grey respectively, (c) Graph-based clusters. (d)-(g) UMAP dimensionality reduction plots of the mammary organoid cells showing the motif accessibility for the indicated TF. (h) GATA1 motif enrichment in sc-ATAC-Seq peaks from Dox vs Ethanol-treated cells. Wilcoxon rank-sum test. (i) Volcano plot of the differential sc-ATAC-Seq peaks from Dox vs Ethanol-treated cells. (j) TF Motif enrichment in the up (red) or down (blue) regulated ATAC-Seq peaks in Dox vs Ethanol-treated cells.

**Supplementary Figure S11:**
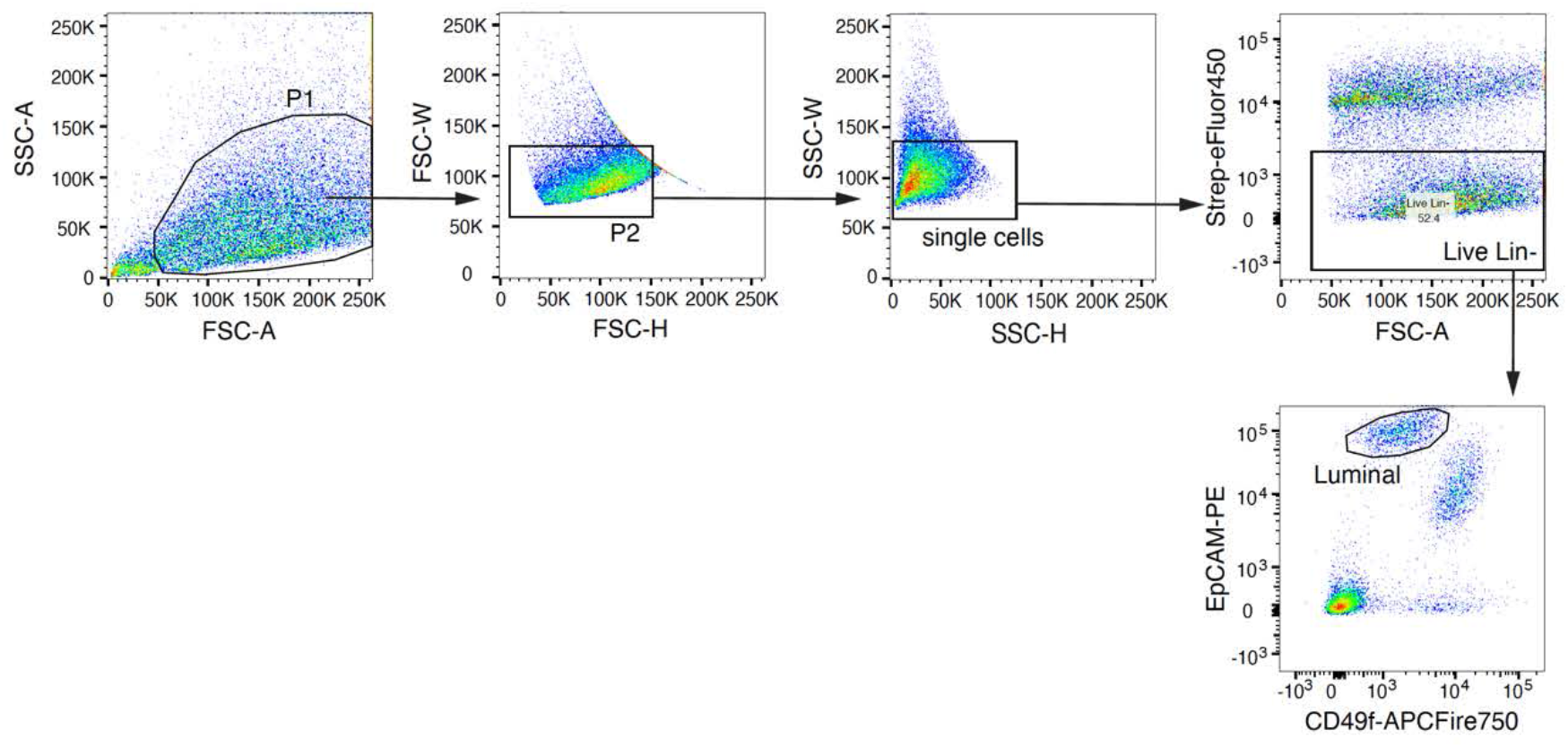
sorting strategy used in Figure 6a to isolate mammary luminal cells from wild type mice for CUT and RUN.

## Notes

### Competing Interest Statement

The authors have declared no competing interest.

