## Supplementary table 2 for "TRPS1 maintains luminal progenitors in the mammary gland by repressing SRF/MRTF activity"

Table S2: Luminal_progenitor_gene_set

Gene_Name

2310002F09Rik

Cd14

Fth1

Celsr2

Pik3r1

Xdh

Golim4

Ccrl2

Stat1

Lrg1

Itga8

Lipa

Mgst1

Ltf

Apobec3

Cd81

Plscr2

Ifngr1

Samhd1

Nqo1

Ndufa4

Nfib

Irs1

R74862

Clic6

Ifit3b

Scd1

Slpi

Xbp1

Crispld2

Dbi

Slc5a8

Nedd9

Dock1

Tgfb3

Crabp2

Map3k1

Prdx1

Irgm2

Taldo1

Slc29a3

4933405E24Rik

Rtp4

Vps28

Tkt

Cep85l

Itpr2

Stat3

Deptor

Plet1os

Plet1

Tubb4b

Pgp

Arrdc3

Pbx1

Ifit3

Plb1

Ifit1

Cited4

Igsf3

Sub1

Uqcrc1

Uqcrc1

Socs5

Mpzl1

Slc16a2

Tcim

Plpp3

Kit

Traf5

Traf5

Lgals3bp

Lcn2

C3

Gm10575

Nhsl1

Maml2

Shisa5

Pgpep1

Thbs1

Ncoa7

Ramp1

Plin2

Phf11d

Igfbp5

Pde4b

Ier2

Fam234a

Gstm1

Mrtfb

Cracr2b

Rb1

Ephx1

Aldoc

Csn3

1700066N21Rik

Hsd17b12

Tcn2

Rab5if

Pitpnc1

Mrpl28

Vcp

Sectm1b

Il10rb

Ctsh

Cp

Id2

Hey1

Trf

Hnrnpm

Atp2b1

P2rx4

Zc2hc1a

Rnd3

Calm1

Idh1

1700022I11Rik

Lypd3

Atp6v1b2

Ucp2

Pmpca

Aldh2

Met

Oasl2

Gstp1

Gstp1

Efna5

Hdac3

Sorbs2

Tcf7l2

Plscr1

Lgals9

Arhgef3

Fxyd3

Cebpb

Mydgf

Ifi35

Efhd2

Wfdc18

Rhoj

Ndufa9

Me1

Ywhab

Clmn

Ifih1

Anpep

Pdgfc

Cyp2d22

Efcab14

Elf5

Trps1

B4galnt1

Ogfrl1

Plekhb1

Osmr

Klk10

Chil1

Parp9

Psap

Lgi4

Slc35a4

Hacd3

Polr3k

Macrod2

Mfsd6

Gm2a

Mrps14

Rab17

Isoc1

Far1

Pgd

Rab4a

Etfdh

Tmem189

Mfge8

Muc1

Coa3

0610040J01Rik

Lurap1l

Mgat2

Rab3d

Vdr

Cd44

Nfix

Acat1

Gstt3

Phlda1

Veph1

Ptx3

Cx3cl1

Gna14

Pla2g4a

Ano3

Muc15

Gas6

Chpt1

Heg1

Cat

Wnt5b

Amotl1

Hebp2

Enpp3

Tmem255b

Gdpd1

1700025G04Rik

9330175M20Rik

Flrt3

Cavin2

Snorc

Aldh1a3
