## Supplementary table 1 for "TRPS1 maintains luminal progenitors in the mammary gland by repressing SRF/MRTF activity"

Table S1: Luminal_mature_gene_set

Gene_Name

Ptn

Cited1

Fxyd2

Tspan13

Qsox1

Ifrd1

Arg2

Slc7a2

Gadd45g

Abhd2

Ktn1

Gipc2

Lcp1

Prlr

Dnaja4

Cystm1

Eif1

Phyh

H1f2

Mt2

Cd24a

Cxcl15

Glul

Hk2

Malat1

Cd164

Dnajb1

H2bc4

Hnrnpc

Tmem56

9530026P05Rik

Ubb

Nek7

Mt1

Tnfrsf21

Gadd45b

Atf4

Cox4i1

4930523C07Rik

Arf6

Tmem159

Tmem132d

Snapc1

Hif1a

Tmed3

Ssh2

Plk2

Btf3

Mtmr7

Ddit3

Hsp90aa1

AW112010

Eif4b

Nr1d1

Gna13

Alcam

Eif4a2

Pgr

Ppp1r15a

Gadd45a

Naca

Atf3

Cdo1

Bhlhe40

Arg1

Gfpt2

Rab31

E230016M11Rik

Tsc22d1

Bnip3

Thra

Etl4

0610040F04Rik

Svil

Phactr4

Itgav

Rps27

Cox7a2l

Rps27rt

Grina

Gde1

Krt18

Itih5

Wfdc2

Adam10

Zc3h15

Sgms2

Sgms2

Rpl18a

Cd200

Ptp4a1

Ptp4a1

Hsph1

Fau

Ctnnbl1

Cav2

Ablim1

Ctla4

Ctla4

Perp

Eyelinc18

Krt7

Flt3l

Rpl13a

Rsrc2

Hs6st3

Etf1

Mboat1

Vmp1

Ffar4

Prom1

Aldh18a1

Rpl5

Jun

Hspe1

G3bp2

Rps18

Cops2

Ndel1

Hp

Klf5

Osbpl10

Sh3d19

Dhx40

Wls

Atg12

Nsd3

Dgat2

Rab5a

Isg20

Lpin2

Eif2s2

Lgmn

Slc22a27

Rbm39

Smarcd1

Eef2

Rps16

Ptpn1

Pfdn5

Elf1

Ovol1

Nfil3

Smagp

Wrap53

Krt19

Dbp

Gcnt2

Pfdn2

Eef1a1

Slc2a1

Tspan3

Ets2

Cdkn1a

Clk1

Commd10

Fbxw11

Tmsb4x

Gm4876

Tgm2

Vdac1

Cnn3

Tes

Nlrp4e

Krt8

Rassf1

Laptm4b

Dipk1a

Cobll1

Tm4sf1

Zfp664

Gpx3

Rpl10a

Ddx50

Il13ra1

Rnf2

Luzp1

Adss

Tuft1

Rps14

Mapre2

Ddit4

Acot1

Meikin

Ocln

Nup50

Oser1

Rpl22

Lgals3

Irf1

Cldn4

Abcb4

Limch1

Elf3

Tfap2b

Ell2

Ecm1

Klf6

Lifr

Prdx6

Bag3

Rpl36

Hagh

Sgms1

Actg1

Ero1l

Abr

Fgb

Clic4

Use1

F3

Rpl32l

9330111N05Rik

Dusp1

R3hdm1

Ctr9

Hpcal1

Srsf11

Nupr1

Zbtb8a

Rpl32

Maea

Adamts6

Pfkfb3

Insig2

Pgam1

Hspa9

Alkbh5

Prdx6b

Pacsin2

Nmbr

1700016L04Rik

Sinhcaf

Hspa8

Large1

Aldh3a2

Hes1

Pik3ip1

Cab39l

Fam162a

Hspb1

Hsp90ab1

Pgm2

Pgm2

Fos

Rhbdd1

Adm

Stc2

Dhx15

Nop58

Higd1a

Bnip3l

Utrn

Cand2

Dsp

Rab11fip1

Tmprss2

Ly6d

Gm6654

Efna1

Prkci

Tgif1

Esr1

Tpi1

Fam104a

Gm13498

Sphk2

Fgg

Sdc1

Fam57b

Bicdl2

Rps26

Uggt1

Rplp1

Map1b

Cdk19

Aldoa

Guk1

Rnf13

Cmtm8

Ccn1

Sntb2

Hspa1b

Rasd1

1810037I17Rik

Nr1d2

Klf9

Cfap77

Cebpg

Hspa1a

Nfyb

9330179D12Rik

Angptl4

Hspd1

Aldoart1

Rdh10

Aldoart2

Pttg1ip

Pgk1

Kdm3a

Anxa2

Fam13a

Ido1

Lratd1

Bsg

Ube2g2

Fam107b

Tor3a

Eif4ebp1

Ubd

Camk2b

Dkkl1

Ets1

Fdft1

Oat

Slc12a2

Ly6a

Otud7b

S100a6

Hspa5

Tmcc3

P4ha2
