## Supplementary table 4 for "TRPS1 maintains luminal progenitors in the mammary gland by repressing SRF/MRTF activity"

Table S10: TRPS1 targets with GO terms related to actin

Gene_Name

Actr1b

Arhgap26

Arhgap28

Arhgap35

Arhgef10l

Atf4

Atf5

Baiap2

Bicdl1

Birc6

C2cd3

Calm3

Calr

Ccnf

Cdc42ep4

Cdk2ap2

Cdk5rap3

Cep55

Cldn3

Cln8

Cttn

Cyld

Dock2

Dstn

Ect2

Epb41

Esrra

Frmd5

Gsk3b

Gsn

Hnrnpc

Hspa2

Incenp

Katnb1

Kbtbd8

Kcnc3

Kif26b

Krt18

Krt8

Lmna

Map3k20

Map4

Msrb1

Mtor

Mvb12a

Myh10

Myh9

Myom2

Ndrg1

Nr3c1

Nubp2

Nudcd2

Pkn2

Poc5

Ppp2ca

Psmd10

Ptp4a1

Rab5a

Rassf7

Rhoa

Rhov

Rtraf

Sbds

Shroom3

Ska2

Slc9a3r1

Son

Specc1

Synpo

Tacc2

Tax1bp3

Tbccd1

Tedc1

Terf1

Tnks1bp1

Togaram1

Tpm1

Ttbk2

Ttll5

Tubb4b

Tubb5

Vcl

Wdpcp

Ypel5
