## Supplementary table 5 for "TRPS1 maintains luminal progenitors in the mammary gland by repressing SRF/MRTF activity"

**Supplementary Table 2: Reagents**

| **REAGENT** | **Source** | **Identifier** | **Concentration** |
| --- | --- | --- | --- |
| **Antibodies** | | |  |
| Biotin anti-TER-119, Clone:TER-119 | ThermoFisher Scientific | 13-5921-82 | 1:50 |
| Biotin anti-CD31, Clone:390 | ThermoFisher Scientific | 13-0311-82 | 1: 50 |
| Biotin anti-CD45, Clone:30-F11 | ThermoFisher Scientific | 13-0451-82 | 1:50 |
| PE-anti mouse EpCAM, Clone:1B7 | Biolegend | 12-9326-42 | 1: 100 |
| APC/Fire^TM^750 anti-human/mouse CD49f antibody, Clone: 93 | Biolegend | 313631 | 1:100 |
| APC anti-mouse CD117 (c-kit)- Clone:2B8 | Biolegend | 105812 | 1: 100 |
| Streptavidin-eFluor450 | ThermoFisher Scientific | 48-4317-82 | 1:200 |
| PE-Vio 770 Anti-mouse CD14 | Miltenyi | 130-115-560 | 1:100 |
| TotalSeq™-A0449 anti-mouse CD326 (Ep-CAM) Antibody | Biolegend | 118237 | 0,5 μg |
| TotalSeq™-A0070 anti-human/mouse CD49f Antibody: | Biolegend | 313633 | 0,5 μg |
| TotalSeq™-A0238 Rat IgG 2α, λ Isotype control | Biolegend | 400571 | 0,5 μg |
| Anti-TRPS1 | Abcam | ab209664 | 1:200 |
| Anti-mouse Krt8 | biolegend | 904804 | 1:500 |
| Anti-mouse Krt8 | Developmental Studies Hybridoma Bank | TROMA-I-s | 2 ug/mL |
| Anti-tGFP | OriGene | TA150041 | 1:400 |
| AF488 Anti-mouse IgG | ThermoFisher Scientific |  | 1:400 |
| AF546 anti-Rabbit IgG | ThermoFisher Scientific |  | 1:400 |
| AF647 anti-Rat IgG | ThermoFisher Scientific |  | 1:400 |
| Anti-Vinculin (hVIN-1) | Sigma-Aldrich | #V9131 | 1:10000 |
| Anti-SRF (D71A9) XP® | Cell signaling | 5147T | 1:100 |
| Anti-mouse IgG-HRP | Santa Cruz | sc-2314 | 1:5000 |
| Anti-rabbit IgG-HRP | Santa Cruz | sc-2313 | 1:5000 |
| **Chemicals, Peptides, and Recombinant Proteins** | | | |
| Gibco Fetal Bovine Serum | ThermoFisher Scientific | F7524-500ML |  |
| Leibovitz's L-15 Medium (500 mL) | ThermoFisher Scientific | 11415064 |  |
| DMEM F-12 Glutamax | ThermoFisher Scientific | 12634028 |  |
| B-27® Supplement (50X) | ThermoFisher Scientific | 17504044 |  |
| N-2 Supplement (100X) | ThermoFisher Scientific | 17502048 |  |
| Murine Noggin, 100µg | Peprotech | 250-38 |  |
| Recombinant Mouse Neuregulin-1/NRG1 Protein, CF, 50µg | Peprotech | 9875-NR-050 |  |
| Leibovitz's L-15 Medium (500 mL) | ThermoFisher Scientific | 11415064 |  |
| Corning® Matrigel® Growth Factor Reduced (GFR) Basement Membrane Matrix, *LDEV-free, | Corning | 354230 |  |
| TrypLE™ Express Enzyme (1X) | ThermoFisher Scientific | 12604013 |  |
| Cultrex Organoid Harvesting Solution | R&D systems | 3700-100-01 |  |
| RPMI 1640 (+GlutaMax) | ThermoFisher Scientific | 21875091 |  |
| EpiCult-B Mouse Medium Kit | STEMCELL Technologies | 5610 |  |
| SYTOX Blue dead cell stain | ThermoFisher Scientific | S34857 |  |
| Giemsa stain | Sigma | GS500 |  |
| **Critical Commercial Assays** | | | |
| innuMIX qPCR DSGreen Standard | Analytik Jena | 845-AS-1300200 |  |
| RNeasy® Micro kit | Qiagen | 74004 |  |
| NEBNext® Ultra RNA Library Prep kit for Illumina | NEB | E7530L |  |
| Chromium Single Cell 3′ Library Construction Kit v3 | 10x Genomics | 1000078 |  |
| Chromium Next GEM Single Cell ATAC Library & Gel Bead Kit v1.1 | 10x Genomics | 1000176 |  |
| MinElute PCR Purification Kit | Qiagen | 28006 |  |
| Agencourt AMPure XP | Beckman Coulter | A63881 |  |
| NEBNext® High-Fidelity 2X PCR Master Mix | NEB | M0541L |  |
| Passive Lysis Buffer | Promega | E194A |  |
| Gentle Collagenase/Hyaluronidase 10X | STEMCELL Technologies | 7919 |  |
