## Supplementary table 6 for "TRPS1 maintains luminal progenitors in the mammary gland by repressing SRF/MRTF activity"

**Supplementary Table: Oligonucleotides**

| **Primer / oligo** | **Sequence** |
| --- | --- |
| **Sc-Seq** | |
| RPI1 | CAAGCAGAAGACGGCATACGAGATCGTGATGTGACTGGAGTTCCTTGGCACCCGAGAATTCCA |
| RPI2 | CAAGCAGAAGACGGCATACGAGATACATCGGTGACTGGAGTTCCTTGGCACCCGAGAATTCCA |
| 10X Genomics SI | AATGATACGGCGACCACCGAGATCTACACTCTTTCCCTACACGACGCTC |
| **Genotyping** | |
| **shRNA genotyping** | |
| shTrps1#1_Fw | AAGCCACAGATGTATCTTCTAATAA |
| shTrps1#2_Fw | AAGCCACAGATGTATTCGTATTTAC |
| shRen_Fw | AAGCCACAGATGTATAGATAAGCAT |
| Col1a1_Rev45 | CACCCTGAAAACTTTGCCCC |
| **Col1a1 genotyping** | |
| Col1a1_Fw | AATCATCCCAGGTGCACAGCATTGCG |
| Col1a1_Rv | CTTTGAGGGCTCATGAACCTCCCAGG |
| SAdpA_Rv2 | AAGACCGCGAAGAGTTTGTC |
| **CAG-lsl-RIK genotyping** | |
| rtTA_Rv2 | CGCTTGTTCTTCACGTGCGA |
| rtTA_Fw (lsl) | AAAAACTCCCACACCTCCC |
| **Rosa26 genotyping** | |
| Rosa_D | TCAGTAAGGGAGCTGCAGTGG |
| Rosa_B | GCGAAGAGTTTGTCCTCAACC |
| Rosa_C | GGAGCGGGAGAAATGGATATG |
| **K8-CreER genotyping** | |
| oIMR7338 | CTAGGCCACAGAATTGAAAGATCT |
| oIMR7339 | GTAGGTGGAAATTCTAGCATCATCC |
| oIMR1084 | GCGGTCTGGCAGTAAAAACTATC |
| oIMR1085 | GTGAAACAGCATTGCTGTCACTT |
| **CAG-rtTA3 genotyping** | |
| SApA_For1 | CTGCTGTCCATTCCTTATTC |
| CH8_Rev2 | CGAAACTCTGGTTGACATG |
| CH8_For1 | TGCCTATCATGTTGTCAAA |
| **qRT-PCR** | |
| mTrps1_Fw | GGTACAGAGGCCACCAGTTAT |
| mTrps1_Rev | GGCTCTCCTTCTACACTTTTGG |
| mAreg_Fw | GGTCTTAGGCTCAGGCCATTA |
| mAreg_Rev | CGCTTATGGTGGAAACCTCTC |
| mCald1_Fw | ATGGTAGAGGAGAAAACACCAGA |
| mCald1_Rev | CCATCCCCTTCTATTTTGGACTC |
| mb2M_1_Fw | AGCCGAACATACTGAACTGCTACG |
| mb2M_1_Rev | CGGCCATACTGTCATGCTTAACTC |
| mActa2_Fw | GTCCCAGACATCAGGGAGTAA |
| mActa2_Rev | TCGGATACTTCAGCGTCAGGA |
| mMyl9_Fw | ACAGCGCCGAGGACTTTTC |
| mMyl9_Rev | AGACATTGGACGTAGCCCTCT |
| mcKit_Fw | GCCTGACGTGCATTGATCC |
| mcKit_Rev | AGTGGCCTCGGCTTTTTCC |
| mElf5_Fw | ATGTTGGACTCCGTAACCCAT |
| mElf5_Rev | GCAGGGTAGTAGTCTTCATTGCT |
| mCd14_Fw | ACTTCTCAGATCCGAAGCCAG |
| mCd14_Rev | CCGCCGTACAATTCCACAT |
| mCtgf_Fw | CAGCATGGACGTTCGTCTG |
| mCtgf_Rev | AACCACGGTTTGGTCCTTGG |
| **shRNAs** | |
| shRen | AAGGTATATTGCTGTTGACAGTGAGCGCAGGAATTATAATGCTTATCTATAGTGAAGCCACA  GATGTATAGATAAGCATTATAATTCCTATGCCTACTGCCTCGGACTTCAAGGGGCTA |
| shTrps1#1 | AAGGTATATTGCTGTTGACAGTGAGCGAGGCGAGCAGATTATTAGAAGATAGTGAAGCCACA  GATGTATCTTCTAATAATCTGCTCGCCGTGCCTACTGCCTCGGACTTCAAGGGGCTA |
| shTrps1#2 | AAGGTATATTGCTGTTGACAGTGAGCGCCCGAGCCTGAGTAAATACGAATAGTGAAGCCACA  GATGTATTCGTATTTACTCAGGCTCGGATGCCTACTGCCTCGGACTTCAAGGGGCTA |
