## Supplementary methods for "TRPS1 maintains luminal progenitors in the mammary gland by repressing SRF/MRTF activity"

**SI Appendix**

*Mice*

Animal experiments were approved by the state government of Thuringia under the animal experiment license FLI-17-004 and FLI-17-017. CAG-lsl-RIK, TRE-tGFP-shTrps1#1, shTrps1#2 or shRenilla [1, 2], and CAG-rtTA3 mice lines were obtained from the Mirimus Company (NY, USA). The shRNA sequences can be found in Supplementary Table 6. To obtain shTrps1 experimental animals and shRen control littermates, homozygous mice carrying both an shTrps1 and an shRen cassette were first generated by breeding CAG-lsl-RIK, TRE-tGFP-shTrps1#1, shTrps1#2 with CAG-lsl-RIK, TRE-tGFP-shRenilla animals. And the resulting progeny was then bred with animals of the CAG-rtTA3 line or the K8-CreER line (C57BL/6J background) (Supplementary Figure S1). Breedings were carried out at the Leibniz Insitute on Aging – Fritz Lipmann institute e.V. (FLI). Wild type mice C57BL/6J or C57BL/6JRj were obtained from The Jackson Laboratory and the Janvier Labs and also bred in house. Mice were used for experiments within an age range of 12 to 20 weeks. The mice were kept in individually ventilated cages (IVCs) under Specific Pathogen-Free (SPF) conditions with a 12 h/12 h dark/light cycle at a temperature of 20°C and a relative humidity of 55% according to the directives of the 2010/63/EU and GV SOLAS. To induce shRNA expression, the animals were fed with doxycycline-supplemented food obtained from the ssniff-Spezialdiäten company (Germany).

*Phenotypic analyses at the German Mouse Clinic*

Mouse phenotyping tests were performed following standard procedures as described before [3, 4](www. mouseclinic.de) and approved by the responsible authority of the district government of Upper Bavaria. Experimental groups were assigned according to the genotype of the animals. The selection of the mice for testing was balanced; control and mutants were measured alternately. Most of the tests were not conducted in blinded conditions because the results were recorded directly by the machines and, therefore, not influenceable by the examiner. The experiment was conducted in blinded conditions whenever there could have been an influence from the investigator. All the procedures are described in SOPs. Metadata for each data point was recorded throughout the measurements, and the influence of this metadata was monitored over time.

X-ray and DXA were performed in an UltraFocus DXA system (Faxitron Bioptics) with automatic exposure control. Histopathological analyses were carried out using H&E stainings of formalin-fixed and paraffin-embedded sections (3 µm) as described in [4] The slides were scanned in a Hamamatsu NanoZoomer 2.0-HT digital scanner and analyzed with NDP.view2 software (Hamamatsu Photonics).

Parameters related to heart function were generated by transthoracic echocardiography (TTE) and electrocardiography (ECG) performed at 12 - weeks-old mice. High-throughput ECG and TTE recordings were performed in conscious mice to assess the electrical conduction system, the morphology and functionality of the heart [5]. Briefly, left ventricular function was evaluated with TTE using a Vevo 3100 Imaging System (Visual Sonics). ECGs were recorded by ECGenie (Mouse Specifics Inc., Boston, MA) and analyzed using e-Mouse software (Mouse Specifics Inc.). High-level details of these procedures have been described [6, 7] and can be accessed publicly:

<https://www.mousephenotype.org/impress/ProcedureInfo?action=list&procID=126>.

If not stated otherwise, data generated by the German Mouse Clinic was analyzed using R (Version 3.4.4). Tests for genotype effects were made by Wilcoxon rank sum test, linear models, or ANOVA and posthoc tests depending on the assumed distribution of the parameter and the questions addressed to the data. A p-value <0.05 has been used as level of significance; a correction for multiple testing has not been performed.

*Isolation of mammary epithelial cells and flow cytometry*

The 4^th^ mouse mammary gland pair was isolated and digested for 15 h in DMEM/F12 medium supplemented with Hyaluronidase/collagenase (STEMCELL Technologies). Tissue lysates were centrifuged, and cell pellets were resuspended in ACK lysis Buffer (150 mM NH_4_Cl, 1M KHCO3, 100 μM EDTA) to eliminate the red blood cells. Cells were then treated with trypsin 0,25 % for 2 min, washed in HF Buffer (Hank’s Balanced Salt Solution + 2% FBS) and treated with a solution containing 5 mg/mL dispase (Merck) and 0.2 mg/mL DNAse (Merck) for 1 min. Cells were washed again in HF buffer, filtered through a 40 μm cell strainer counted and transferred to PBS 2% FBS. Cells were then incubated with a mixture of biotin-conjugated lineage antibodies (against TER-119, CD31 and CD45, Table S5). After washing in PBS 2% FBS, the cells were incubated with fluorophore conjugated antibodies (PE-Anti-mouse EpCAM, APCFire750- anti-mouse CD49f and APC-anti -ckit, PEvio770-anti mouse CD14, See Table S5) and strepatavidin-eFluor450 (Table S5).

For flow cytometric analysis and subsequent sorting, stained cells were washed twice and resuspended in PBS 2% FBS. The cell suspension was then filtered through (40 µm) and 1 μM SYTOX Blue dead cell stain (Thermo Fisher Scientific) was added. Fluorescence-activated sorting panels were set up using the automatic compensation feature of the FACS Diva software (BD Biosiences) according to the manufacturer’s instructions with unstained cells or Ultracomp eBeadsTM (Thermo Fisher Scientific) as single-stained compensation controls (see the eBeadsTM manual for the protocol). Cells were sorted into low binding round bottom 96-well plates in medium using the BD FACSAria Fusion cell sorter (BD Biosciences) and were stored on ice until further processing.

*CITE-Seq*

For sc-Seq of the mammary gland cell populations, the lymph node was removed prior to mammary gland cell isolation (see *Isolation of mammary epithelial cells and flow cytometry*) and blood lineage were depleted by incubating the cells with a mixture of biotin-conjugated lineage antibodies (Thermo Fisher Scientific) (against TER-119, CD31 and CD45), washing in PBS 2% FBS and incubating with anti-Biotin microbeads (ref). The cells were then washed in PBS 2% FBS, resuspended in PBS and loaded on a LS MACS column (Miltenyi). Cells transferred to PBS + BSA 2% + Tween 20 0,01% and incubated with Fc blocking reagent 10 min at 4°C. 0.5 ug of each Total Seq antibodies (anti-EpCAM and anti-CD49f, Biolegend) was added to the cell suspension and the samples were incubated 30 min at 4°C. Cells were washed in PBS + BSA 2% + Tween 20 0,01% and cells were counted with an automated cell counter (EVE, NanoEnTek) and adjusted to a concentration of 1000 cells/ µL. Cells were subjected to the Chromium single cell 3’ assay v3 (10X Genomics) as recommended by the manufacturer. After cDNA amplification, ADT-derived cDNAs and mRNA-derived cDNAs were separated based on their size using 0.6x AMPure XP Beads (Beckman Coulter). The mRNA-derived cDNAs contained in the bead fraction were further processed following the standard 10X Genomics protocol in order to generate single-cell (sc)RNA libraries. The ADT-derived cDNAs contained in the supernatant were further purified as follows: 1.4x beads were added to obtain a final AMPure beads concentration of 2X and samples were incubated 10 min at RT. Supernatant was discarded and beads were washed with Ethanol 80%, air-dried and resuspended in 50 µL water. The same procedure was repeated once. Beads were washed twice with Ethanol 80% and air-dried. The purified ADTs were subsequently eluted in 45 µL water at RT for 5 min. To generate the ADT sequencing library, 45 µL ADTs were used as template in a 100 µL PCR reaction with the NEBNext^®^ High Fidelity Master Mix (NEB), a Truseq small RNA RPIx (containing i7 index) primer and the 10X Genomics SI-PCR primer (see primer table). The PCR products were incubated with 1.6x AMPure beads at RT for 5 min. After washing the beads twice with Ethanol 80%, the ADT library was eluted in 30 µL water. The libraries were quantified using an Agilent Tapestation 4200. The scRNA-Seq libraries and the ADT libraries were sequenced together on an Illumina NextSeq500 platform using a 75 cycle High Output v2.5 kit. The following sequencing cycles were performed: R1 (10x barcode + UMI): 26bp; R2 (cDNA): 53 bp; i7 index: 8 bp. Extraction of FastQ files was done using bcl2fastq v2.20.0.422 (Illumina).

*Mammary gland organoids isolation and cultivation*

The 4th pair of mouse mammary glands was isolated and finely chopped with a scalpel. The tissue was transferred to a tube adapted to the Octo tissue dissociator from Miltenyi containing 2.5 mL of Leibovitz's L-15 Medium (Thermo Fisher Scientific) supplemented with 3 mg/mL Collagenase A and 1.5 mg/mL Trypsin. Samples were incubated in the Octo tissue dissociator 1h at 37°C, 100 rpm. Samples were then centrifuged; the cell pellet was resuspended in DMEM/F12 medium and filtered through a 70 µm cell strainer. The samples were centrifuged again, and the pellet was resuspended in 50 µL of Matrigel (Corning). 10 µL of organoid suspension was dropped in 5 wells of a pre-warmed 48-well plate. The plate was placed at 37°C to allow the Matrigel drops to solidify and 250 µL of Organoid medium (DMEM/F12 + Glutamax with addition of 1X N-2 Supplement, 1X B27 Supplement, 100 ng/mL Neuregulin, 100 ng/mL Noggin and 100 ng/mL R-spondin) was added in each well. Medium was exchanged twice a week and every 2 to 3 weeks, the organoids were passaged. For passaging, the matrigel domes were washed with PBS and depolymerized in 250 µL CultRexTM organoid harvesting solution (R&D systems) on ice for 1-2h. The organoid suspension from several wells was combined in one centrifugation tube and washed in cold PBS. The organoids were partially dissociated in TrypLE Express Enzyme (Thermo Fisher Scientific) for 2 to 5 min at 37°C. After washing with PBS, the organoids were resuspended in matrigel and seeded again in a new 48-well plate.

*Colony forming assay*

Luminal or luminal progenitor cells were isolated from mouse mammary glands and sorted in 96-well plate filled with EpiCULT medium (STEMCELL Technologies) supplemented with 5% FBS, 10 ng/mL EGF, 10 ng/mL FGF and 4 µg/mL heparin. 500 luminal cells or 200 luminal progenitors were then transferred to a 48-well plate previously filled with 10,000 NIH3T3 irradiated feeder cells per well and incubated at 37°C, 5% CO_2_. 24h later the medium was exchanged for EpiCULT medium supplemented, 10 ng/mL EGF, 10 ng/mL FGF and 4 µg/mL heparin and without FBS. When needed, doxycycline (10 µg/mL) or Ethanol was added (1:1000) to the culture medium. Cells were incubated for 7 days and medium was exchanged every 2 days. Colonies that ultimately formed were fixed for 1 min in methanol and stained in 10% Giemsa staining solution for 20 min.

*Western blotting*

Cells were lysed in RIPA buffer (50 mM Hepes pH 7.9, 140 mM NaCl, 1 mM EDTA, 1% Triton X-100, 0.1% Na-deoxycholate, 0.1% SDS) containing sodium pyrophosphate and protease inhibitor cocktail (Sigma). Lysates were cleared by centrifugation, separated on 8% Bis-Tris gels and transferred to a PVDF membrane (Millipore). Membranes were blocked with 5% skim milk powder in TBS, probed with primary antibodies diluted in 5% BSA in TBS and finally incubated with the appropriate horseradish peroxidase-coupled secondary antibodies. Visualization was performed using chemiluminescence HRP substrate (Immobilon Western, Millipore).

*RNA-Sequencing*

For all RNA-Sequencing samples, three biological replicates per condition were analyzed. Total RNA was extracted using RNeasy® Micro Kit (Qiagen) with on-column DNaseI (Qiagen) digestion. RNA integrity (all processed samples had a RIN>8) was verified with the Agilent Bioanalyzer 2100 automated electrophoresis system (Agilent Technologies). mRNA (at least 10 ng) was isolated using the NEBNext® Poly(A) mRNA Magnetic Isolation Module (NEB) and library preparation was conducted with the NEBNext® Ultra RNA Library Prep Kit for Illumina (NEB) with Dual Index Primers (NEBNext® Multiplex Oligos for Illumina, NEB) or with NEBNext® Single Cell/Low Input RNA library kit (NEB) for low number of cells following the manufacturer’s description. Cycles for amplification of the cDNA were determined by qRT-PCR. Libraries were quantified with the Agilent 2100 Bioanalyzer automated electrophoresis system (Agilent Technologies) and subjected to 51 bp single-end Illumina Sequencing on a HiSeq 2500 in high-output mode. Reads were extracted in FastQ format using bcl2fastq v1.8.4 (Illumina).

*RNA-Sequencing analysis*

Adapter removal, size selection (reads > 25 nt) and quality filtering (Phred score > 43) of FASTQ files was performed with cutadapt (http://cutadapt.readthedocs.io/en/stable/guide.html#). Reads were then aligned to the mouse genome (mm10) using bowtie2 (v2.2.9) using default settings. Differential gene expression analysis was performed with edgeR (v3.26.8) using default parameters. PCA analysis was performed using DESeq2 on the 500 most variable genes.

*qRT-PCR*

RNA was extracted with peqGOLD TriFast Reagent (Peqlab). First-strand cDNA synthesis was performed using M-MLV Reverse Transcriptase (Promega) and random hexamers (Sigma) according to standard procedures. PCR reaction was performed in technical triplicates using Innumix SybrGreen Mix (Analytik Jena). Gene expression was analyzed with a StepOnePlus™ Real-Time PCR System (Thermo Fisher Scientific). The expression values were normalized to *B2m* as housekeeping gene using the ddCt method. The primer sequences are listed in Supplementary Table 6.

*CUT&RUN*

Briefly, for each CUT and RUN reaction 200,000 sorted mouse luminal cells were washed, resuspended in 100 μl wash buffer (20 mM HEPES, pH7.5, 150 mM NaCl, 0.5 mM Spermidine) and bound to 10 μl activated concanavalin A magnetic beads for 10 min at room temperature. Bead-bound cells were then incubated in 100 µl antibody buffer (wash buffer + 0.01% digitonin and 2 mM EDTA) with the desired antibody (1:100) overnight at 4°C. As negative control an IgG rabbit antibody was used. Beads were washed in digitonin wash buffer (wash buffer + 0.01% digitonin) and incubated 1h at 4°C with 1 μg/mL protein A/G Micrococcal Nuclease fusion protein (pA/G MNase). After 3 washing steps in digitonin wash buffer, beads were rinsed with Low salt buffer (20 mM HEPES, pH7.5, 0.5 mM Spermidine, 0.01% digitonine) and placed in 200 µl incubation buffer (20 mM HEPES, pH7.5, 10 mM CaCl_2_, 0.01% digitonin) at 0°C to initiate cleavage. After 30 min, reactions were stopped by adding 200 µl STOP buffer (170 mM NaCl, 20 mM EGTA, 0.01% digitonin, 50 µg/ml RNAse A) and samples were incubated 30 min at 37°C to digest the RNA and release the DNA fragments. The samples were then treated with proteinase K for 1h at 50°C and the DNA was purified using Phenol/Chloroform/Isoamyl alcohol. After precipitation with glycogen and Ethanol, the DNA pellet was resuspended in 0.1 X TE and used for DNA library generation with the NEBNext^®^ Ultra™ II DNA Library Prep Kit for Illumina^®^ (New England Biolabs) according to the manufacturer’s recommendations. Adaptor ligation was performed with 1:25 diluted adaptor and 15 cycles were used for library amplification.

*scATAC-Seq*

Cells were isolated from mammary gland organoids from three different TRPS1 depletion mice of the sh*Trps1*#2 line. Prior to cell isolation, organoids were treated for 7 days with doxycycline (1:1000) or with Ethanol as control. Organoids were harvested in CultRexTM organoid harvesting solution (R&D systems), washed with PBS and dissociated in TrypLE Express Enzyme (Thermo Fisher Scientific) for 3 min at 37°C, washed again with cold PBS. Cells debris were eliminated using debris removal solution according to the manufacturer’s recommendation (Miltenyi). Cells were then resuspended in PBS 0.04% BSA and counted. For each sample (Dox treated or Ethanol control) 20,000 cells from each animal were mixed and cells were transferred into a round-bottom 96 well plate (60,000 cells in total per sample). Nuclei were isolated by lysing the cells with chilled lysis buffer (10 mM Tris-HCl pH 7.4, 10 mM NaCl, 3 mM MgCl2, 0.1% Tween 20, 0.1%Nonidet P40, 0.01 % Digitonin, 1 % BSA) on ice for 3 min 30 s. Cells were washed with wash buffer (10 mM Tris-HCl pH 7.4, 10 mM NaCl, 3 mM MgCl2, 0.1% Tween 20, 1 % BSA). Supernatant was discarded and cells were resuspended in 1X Nuclei Buffer (10X Genomics, PN-2000153/ 2000207).

Nuclei suspensions were processed using the 10x Genomics Chromium Controller and Chromium Next GEM Single Cell ATAC Reagent Kits v2 according to the manufacturer’s protocol (CG000496). 7600 nuclei were loaded onto the Chroimium Controller to recover 5000 nuclei for library prepration and sequencing. Total yield and quality of cDNA was assesed on DNA 7500 assay (Agilent 2100 Bioanalyzer). The libraries were pooled and sequenced on Illumina NovaSeq 6000 System in combination with SP 100 cycles v1.5 kit. The following sequencing cycles were performed: R1 (sequencing of interest: 51 bp; R2 (sequencing of interest): 51 bp; i7 index (sample index): 8 bp; i5 index (10X barcode): 16 bp. Extraction of FastQ files was done using bcl2fastq v2.20.0.422 (Illumina).

*scATAC-Seq analysis*

FASTQ files were analyzed by the 10X Genomics cellranger-atac pipeline (v1.2.0). The transcription factor motif accessibility is taken from the cellranger output. ChromVAR was used to cluster the cells based on their accessibility in peaks for all motifs from the JASPAR2016 database comprising 386 transcription factor binding motifs.

*Immunofluorescence on mammary gland tissue sections*

Mammary gland samples were fixed in PFA 4% for 1h at RT rinsed in PBS and Ethanol 50% and embedded in paraffine. Paraffine blocks were cut in 10 µm-thick sections. Sections were blocked for 1h in blocking buffer (1X TBS, 0.3% Triton X-100, 5% Goat serum, Fab Fragments goat anti-mouse) and incubated with a mixture of primary antibody (against TRPS1, GFP and Krt8, see Table S5) overnight at 4°C. Sections were then washed 3 times with 1X TBS and incubated with a mixture of secondary antibodies (AF546-anti-rabbit, AF488-anti-mouse, AF647-anti-rat, see Table S5) for 30 min at RT. After washing with 1X TBS, sections were stained with Hoechst (1:1000), washed again and mounted into Fluoromount medium (Thermo Fisher Scientific). Pictures were taken using an Apotome microscope (Zeiss).

Cryosections were obtained by flash freezing fresh mammary gland tissue in x medium. Frozen blocks were then cut in 10 μm-thick sections. Sections were fixed in PFA 4% for 10 min at RT and stained the same way as the paraffine embedded sections.
